# Structural integrity moderates the cognitive bearing of dopamine clearance in aging

**DOI:** 10.64898/2026.09.29.753541

**Authors:** Amirhossein Esmaeili, Giacomo Preti, Nora Bittner, M Marmaduke Woodman, Svenja Caspers, Meysam Hashemi, Spase Petkoski, Daniele Marinazzo, Viktor Jirsa

## Abstract

Age-related cognitive decline in humans is well established, yet how neurobiological variations influence brain dynamics, and subsequently behavioral performance, remains insufficiently explored. Microscale accounts of cognitive aging revolve around attenuations in dopamine (DA) receptors and transporters (DAT). In contrast, macroscale accounts associate individual declines in cognitive performance with functional brain networks’ reorganization and white-matter tracts’ deterioration. We endeavor to bridge this gap with an amortized dopaminergic aging model (ADAM) which assembles aging brain twins by coupling dopaminergic modulation with structural connectomes to constrain emergent whole-brain dynamics. ADAM thereby augments established DAT decline with personalized estimates, through a reusable and generalizable estimator capable of disentangling neuromodulatory from structural contributions to age-related decline within each individual. To that end, we embed structural connectomes into a low-dimensional space exhibiting high reconstruction fidelity and —pseudonymized— individual identifiability. The spectrum of structural embeddings is subsequently updated with fMRI observations to estimate DA clearance timescales in 3,650 adults aged 36 to 95 years across five independent aging cohorts. Our analysis reveals an age-related elevation of DA clearance timescales. Critically, low performers exhibit elevated clearance in adults with marked structural degradation, and only within sliding age windows centred between 56 and 76 years. This supports a contribution of dopamine to aging, and reveals a structure- and age-conditional association between its clearance and cognitive performance. Our findings suggest that structural connectivity gates the macroscale reach of dopaminergic modulation, thus rendering the gain-control framework of neuromodulation mechanistically explicit across individuals and cohorts. ADAM therefore provides a biologically plausible framework that anchors interindividual variability in cognitive aging to perturbations in the dopaminergic system by learning cohort-scale mappings which can be shared across sites, establishing a transferable testbed for precision neuromodulation and future in-silico pharmacological trials.

## Introduction

To age with strength and independence, so that our later years remain as vibrant as our younger ones, we have to grapple with the usual companion of aging, the mind’s protracted decline. Our current understanding of the aging brain underscores a progressive decline and reorganization of neural subsystems, portrayed through observations ranging from single neurons (i.e. microscale) to whole-brain (i.e. macroscale) networks. Nevertheless, a systemic and mechanistic portrait of the interplay between these subsystems, particularly in relation to personalization of cognitive performance variability, is yet to be sufficiently charted.

Macroscale aging frameworks peer through a coarse-grained yet individually resolvable lens, and often revolve around multimodal perturbations to network connectivity. Pertaining to structural perturbations, older adults have been shown to exhibit declines in interhemispheric connectivity and white matter integrity (Sullivan et al. 2010; Westlye et al. 2010; Puxeddu et al. 2020; Stumme et al. 2022). White matter degradation may also follow a distinct lifespan trajectory (Bethlehem et al. 2022) and bear age- and cognition-related associations beyond those derived from grey matter alone (Cong et al. 2026). Declines in functional connectivity (FC) and in its ongoing reconfiguration (functional connectivity dynamics, FCD) commonly accompany the aforementioned structural degradation, most prominently as reduced homotopic FC and diminished interhemispheric fluidity (i.e. attenuated reconfiguration of FC over time; Hansen et al. (2015); Battaglia et al. (2020); Petkoski et al. (2023)). Crucially, macroscale decline has been shown to track cognition. Subjects exhibiting the steepest functional decline, for instance, also show the sharpest concurrent losses across domains such as processing speed, executive function, and memory (Chan et al. 2014; Stumme et al. 2022). This finding has been corroborated longitudinally, where the loss of functional segregation tracks both attenuating white-matter integrity and cognitive decline within the same individuals (Pedersen et al. 2021).

At the opposite end of said spatiotemporal scales, microscale (i.e. synaptic or neuronal) hypotheses of cognitive aging routinely implicate the dopaminergic (DA) system. A prime example is a comprehensive metaanalysis of positron emission tomography (PET) and single-photon emission computed tomography (SPECT) studies which reports an age-dependent decline in both D1 receptors (D1Rs) and DATs, with dopamine synthesis capacity being essentially preserved (Karrer et al. 2017). Previous inquiries have made strides in bridging these scales, with evidence suggesting that reduced D1R availability accompanies functional over-integration and tracks the concomitant decline in task performance (Pedersen et al. 2023; Korkki et al. 2025). Nevertheless, to the best of our knowledge, no attempt has reconciled them mechanistically in a manner which integrates *transporter-mediated* dopamine clearance with macroscale functional decline and, most importantly, with structural connectivity attrition most prominently observed in interhemispheric connections. The framework of neural gain-control may however serve as the theoretical bedrock for this reconciliation. Within it, neuronal gain is represented by a control parameter which establishes the slope of the sigmoidal activation function relating preto post-synaptic activity, such that attenuated gain flattens that function and renders a unit less responsive to its inputs (Li et al. 2001; Li and Rieckmann 2014). Neuromodulation consequently influences cognition by varying the gain, and thereby the signal-to-noise ratio (SNR) of neuronal input-output functions (Li et al. 2001; Li and Rieckmann 2014). However, transient changes in dopamine are consequential only in the presence of lower tonic levels, both in terms of changes to neural gain, and of cognitive function (Kroener et al. 2009; Cools and D’Esposito 2011).

This baseline inspires our in-silico target in the present work. Within the young adult brain, the abundance of D1 receptors and rapid DAT-mediated clearance (Karrer et al. 2017) are modelled as producing a highcontrast DA modulation which sharply switches between elevated values and a comparatively lower baseline (Dreher and Burnod 2002). In older adults, on the other hand, slow DA unbinding (Hunger et al. 2020) alongside DAT-mediated elongation of clearance timescales may result in sustained excitatory drive (at the loss of phasic modulation found in young adults), with this excess excitation potentially driving regions towards the ceiling of their dynamic spectrum (Thurley et al. 2008; Sun et al. 2005). At the macroscale, regions in proximity to this ceiling lose input responsiveness (Kroener et al. 2009), leading to the loss of representational distinctiveness anticipated by the gain-control framework (Li et al. 2001). Aging degrades the clasp of the structural scaffold on neural dynamics (Zhang et al. 2024), and this reduction of structure-function coupling permits neuromodulation to exert a greater influence over the dynamics. We accordingly posit that prolonged effective clearance becomes consequential for macroscale dynamics, and for cognitive performance, under structural attrition.

Upon initial glance, any attempt to estimate DA clearance timescales in living participants faces a seemingly insurmountable impasse. Diffusion MRI and resting-state fMRI offer whole-brain coverage but are unable to estimate receptor-level or transporter-level kinetic parameters. Conversely, molecular imaging techniques which target such kinetic constants and timescales (PET, SPECT) in practice yield receptor and transporter *availability* under kinetic-model assumptions, and are further constrained by tracer availability, radiation exposure, and cost. These constraints are compounded by the nature of the age trajectories derived from such measurements, which are normative rates pooled cross-sectionally across studies and participants (Karrer et al. 2017). Subject-specific DA clearance timescales and their influence on network-scale connectivity are therefore not directly measurable. Alleviation of these constraints hence necessitates a multimodal generative framework which can plausibly represent clearance dynamics, accommodate the concurrent decline in functional and structural connectomes, and elevate previous virtualization schemes from the scale of subjects towards that of cohorts. Virtual brain twins (VBTs) are at the heart of our proposed generative framework. VBTs are whole-brain models which are personalized to an individual’s neuroimaging (or electrophysiological) data (Hashemi et al. 2025) across myriad in-silico representations, such as epilepsy (Hashemi et al. 2023) and schizophrenia (Preti et al. 2026). Nevertheless, virtual twin formulations have been commonly employed per-subject, and thus cannot be rigorously generalized to population-scale (i.e. normative) trends.

Enter amortized Bayesian inference, a principled approach which allows one to estimate difficult-to-measure model parameters in light of observations. Amortized inference enables one to derive a reusable mapping from the entire set of observables to model parameters once, and to subsequently apply said mapping to new (empirical) observations, bypassing the need for re-training (Hashemi et al. 2023). It can accordingly transform a normative measurement into a personalized one by evaluation of subjects’ neuroimaging data through inherited population-scale priors. Cohort-amortized personalization (Esmaeili et al. 2026), a recently proposed framework for generating and inverting mechanistic whole-brain models at cohort scale, is therefore well-positioned for estimation of virtual twin configurations which are participant-specific yet bounded within their normative source.

The present work formulates a cohort-amortized virtual brain twin (Esmaeili et al. 2026; Hashemi et al. 2025) that explicitly models and subsequently estimates personalized dopaminergic clearance timescales. The characteristic time constant *τ*_clear_ represents (and sets) this timescale, such that larger *τ*_clear_ translate to slower effective transporter-mediated uptake within the model. To that end, we extend a mechanistic neural population model (Gast et al. 2024; Preti et al. 2026; Depannemaecker et al. 2026) incorporating D1-receptor modulation of AMPA synaptic conductance and extracellular dopamine concentration dynamics governed by Michaelis– Menten transporter kinetics. Our amortized dopaminergic aging model (hereafter referred to as ADAM) is constrained to D1 signaling. We confined our inquiry to this pathway as it is linked to cortico-striatal age-related over-integration and poorer memory (Pedersen et al. 2023; Korkki et al. 2025), is more age-sensitive than D2-like receptors (Karrer et al. 2017; Pedersen et al. 2023), and is more prevalent than D2 in primate prefrontal pyramidal neurons (Mueller et al. 2020), where it modulates AMPA-receptor trafficking and neuronal gain (Sun et al. 2005; Thurley et al. 2008). In contrast to previous aging twin formulations (Lavanga et al. 2023), ADAM introduces a biologically plausible neuromodulatory basis on top of the structural axis, wherein the cognitive pertinence of dopaminergic clearance and white matter degradation can be estimated individually.

In the following, we embed structural connectomes across five independent aging cohorts, totalling *N* = 3,650 participants and spanning an age range of 36 to 95 years (Methods, Section 1.1), into two coordinates that preserve reconstruction fidelity while keeping individual participants distinguishable. These coordinates supply a normative range of connectome characteristics across the cohorts which, together with one over the clearance timescales *τ*_clear_, are employed for learning a reusable mapping between them and their corresponding synthetic macroscale imaging summaries (i.e. summary statistics computed from structural connectomes and BOLD time-series). Subsequently, we leverage this mapping to estimate distributions over dopaminergic clearance and connectome coordinates for any participant in light of empirical imaging summaries. Our findings indicate an elevation of *τ*_clear_ with age, corroborating the previously reported age-related decline in DAT availability (Karrer et al. 2017). We finally qualify this elevation with the influence of said timescales on cognition, which we show to be heterogeneous across the aging population, and largely confined to individuals with pronounced structural attrition.

## Results

### Multimodal signatures of brain aging across five independent cohorts

We compiled a cross-cohort multimodal dataset from five independent aging cohorts, spanning a combined age range of 36–95 years: 1000BRAINS (*N* = 841; (Caspers et al. 2014)), HCP-Aging (HCP-A, *N* = 1,208; (Bookheimer et al. 2019)), NKI-Rockland Sample (NKI-RS, *N* = 544; (Nooner et al. 2012)), Cam-CAN (*N* = 511; (Shafto et al. 2014; Taylor et al. 2017)), and OASIS-3 (*N* = 546; (LaMontagne et al. 2019)) (Fig. 1*d*).

**Fig. 1:**
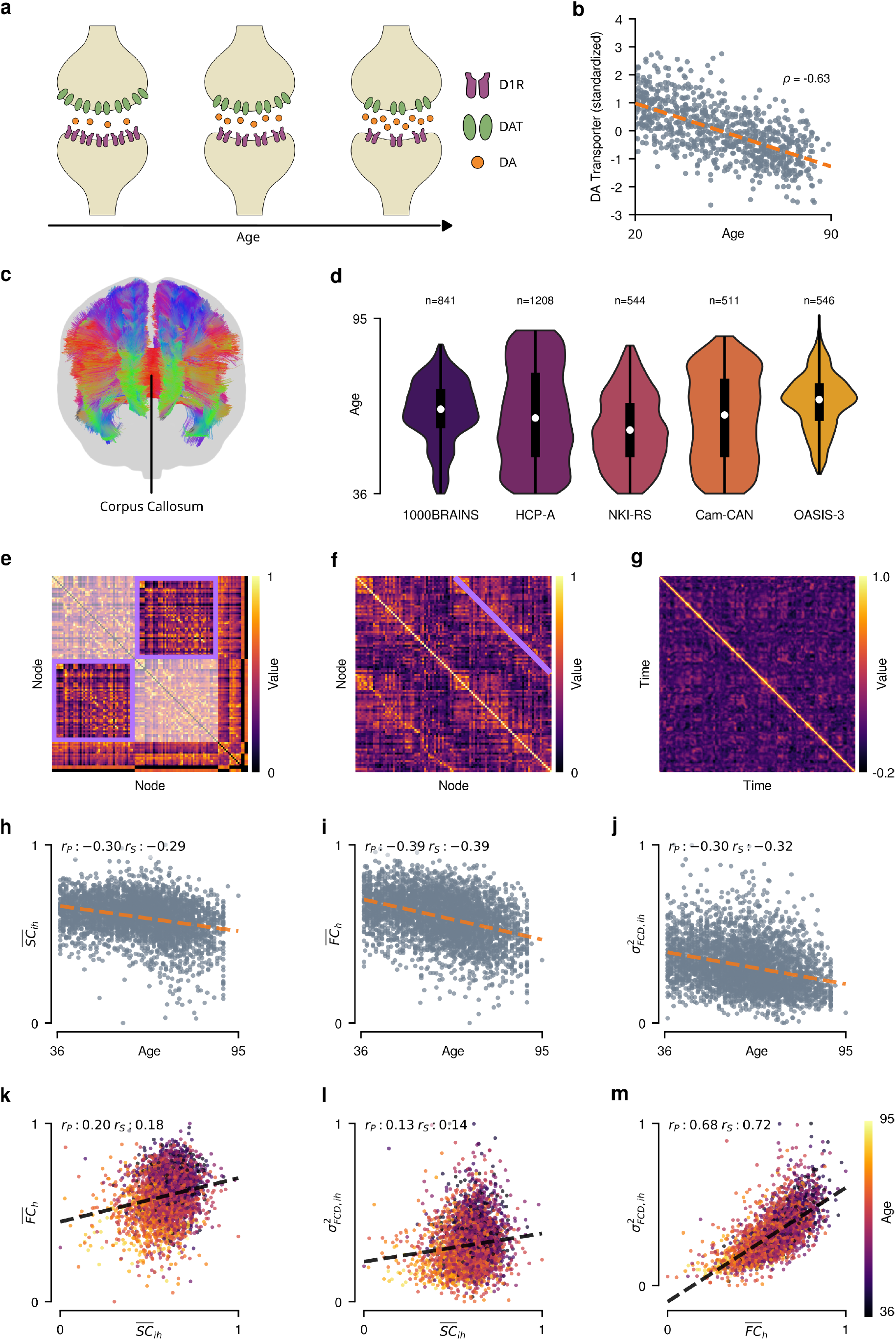
Multimodal empirical observations across five aging cohorts. **a**, Schematic of the dopaminergic synapse across the lifespan, illustrating D1-receptor binding and DAT-mediated reuptake, and highlighting the age-related decline of D1-receptor density and DAT availability. Cleft dopamine is schematised as a time-averaged steady-state extracellular concentration, not a single-release transient. **b**, Standardized DAT availability as a function of age, compiled from the PET/SPECT meta-analysis of Karrer et al. (2017), showing a significant negative association (*ρ* = *−*0.63). **c**, Representative whole-brain tractogram, illustrating corpus callosum as the principal interhemispheric connective substrate. **d**, Age distributions of the five aging cohorts (1000BRAINS, HCP-A, NKI-RS, Cam-CAN, OASIS-3), displayed as combined violin-and-box plots along with the corresponding subject counts. **e**, Representative empirical SC matrix from an individual subject, obtained from probabilistic tractography over the 114-node Schaefer-100 + Harvard-Oxford-14 parcellation, alongside cohort-scaled dopaminergic projection sources (bilateral VTA and SNc); the two interhemispheric quadrants are outlined in violet and the intrahemispheric quadrants overlaid with a translucent white mask, so as to foreground the cross-hemisphere edges that carry 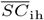. For visual simplicity the overlay is shown over the 100 cortical nodes alone, across which the hemispheres partition evenly. **f**, Representative empirical FC matrix from an individual subject, with homotopic connections highlighted. **g**, Representative empirical interhemispheric FCD matrix from an individual subject. **h–j**, Scatter and regression plots of the three empirical summary features — mean interhemispheric SC 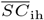, mean homotopic FC 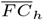, and interhemispheric fluidity 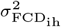, respectively — as a function of age, colour-coded by cohort. All three features display significant age-related decline across the pooled sample (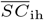: *r* = *−*0.30, *p* = 1.3 *×* 10^*−*74^; 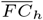 : *r* = *−*0.39, *p* = 8.4 *×* 10^*−*133^; 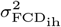 : *r* = *−*0.30, *p* = 4.0 *×* 10^*−*77^). **k–m**, Pairwise scatter and regression plots between the three empirical summary features: 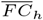 vs. 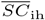 (*r* = 0.20), 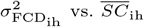 (*r* = 0.13), and 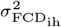 vs. 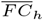 (*r* = 0.68), respectively.

For four out of five cohorts, we derived structural connectomes (SC) from diffusion-weighted imaging and wholebrain probabilistic tractography (Tournier et al. 2019; Behrens et al. 2007) by employing a 114-node parcellation combining the Schaefer 17-network 100-parcel atlas (Schaefer et al. 2018) with 14 Harvard-Oxford subcortical nodes (Frazier et al. 2005; Makris et al. 2006; Goldstein et al. 2007). These were combined with cohort-scaled substantia nigra pars compacta (SNc) and bilateral ventral tegmental area (VTA) weights derived from a template (Fig. 1*c, e*). Similarly parcellated BOLD time-series arrays were computed from resting-state fMRI following a per-cohort denoising sequence (Methods, Sections 1.1.1 and 1.1.3) (Fig. 1*f*). Finally, for each subject, we conducted principal-component analysis on a subset of each cohort’s cognitive performance metrics, the first component of which was employed in the entirety of the study (Stumme et al. 2022) (Methods, Section 1.1.5). First, our results not only confirm previously reported age-related decline of summary statistics within both functional and structural connectomes, but extend it to all five cohorts. We computed all summary statistics over the 100-node cortical subset of the parcellation (circumscribed to personalized connectivity; see Methods Section 1.3.1) with this subset being common to all downstream analyses (Methods, Section 1.4, and Supplementary Section S5., for a comprehensive account of all data features). We observed a significant attenuation of said summaries with advancing age including mean interhemispheric structural connectivity 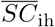 (i.e. average weight of the cross-hemisphere edges of the connectome), mean homotopic functional connectivity 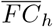 (i.e. average of the FC between each brain region and its contralateral homotope), and interhemispheric FCD variance (hereafter referred to as interhemispheric *fluidity*; *r* = *−*0.30, *r* = *−*0.39, *r* = *−*0.30, with *p* = 1.3 *×* 10^*−*74^, *p* = 8.4 *×* 10^*−*133^, *p* = 4.0 *×* 10^*−*77^, respectively; Fig. 1*h*–*j*). This finding is aligned with age-related attrition of white matter integrity, homotopic FC, and interhemispheric fluidity in the 1000BRAINS cohort as reported by a prior virtual brain twin formulation (Lavanga et al. 2023), and with the wider research on aging connectomes (Puxeddu et al. 2020; Stumme et al. 2022; Petkoski et al. 2023; Mollink et al. 2019; Roland et al. 2017). Interhemispheric fluidity is especially salient here, due to its attenuation reflecting reduced switching between the brain’s non-stationary functional states (Hansen et al. 2015), in addition to serving as a dynamical signature of age-related decline (Lavanga et al. 2023; Petkoski et al. 2023; Battaglia et al. 2020). Aside from the aforementioned age-related trajectories, thinning of interhemispheric white matter was concomitant with attenuation of homotopic functional connectivity and dwindling of dynamical state-transitions 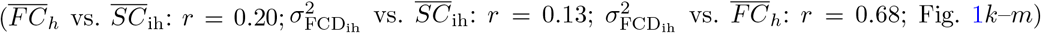 This pattern is suggestive of structural attrition restricting but not unequivocally determining its accompanying functional decline.

Nevertheless, structural decay by itself does not fully account for interindividual variability in cognitive decline (see *Cohort-amortized personalization of DA clearance timescales* below and Supplementary Section S8.), which impels the mechanistic model constructed in the remainder of this investigation. Regardless of the age-association, we implemented cross-cohort harmonization across all data features, to attain comparable baselines of summary statistics across all cohorts, which is a pivotal prerequisite of our cohort-amortization scheme. In the following, we embed this harmonized, cross-cohort signature within a mechanistic amortized dopaminergic model of aging.

### Amortized Dopaminergic Model of Aging

We developed a two-compartment data generative framework so as to model the age-related joint attenuation of functional and structural connectomes. Our implementation couples a cohort-informed structural prior to inform a mechanistic whole-brain dynamical model, which are trained jointly thereafter under a unified amortized network. This coupling yields a joint posterior over *{θ*SC_1_, *θ*_SC2_, *τ*_clear_*}* enabling uncertainty quantification across the structural and neuromodulatory components simultaneously. Figure 2 provides a portrait of this pipeline, from individual components of forward simulation afforded by the whole-brain model (with Fig. 3 portraying their coalescence for conducting cohort-scale amortized inference). Our regional-level model extends the previously derived mean-field estimation of a network of Izhikevich neurons with additive conductance-based excitatory (AMPA) and inhibitory (GABA) synapses (Gast et al. 2024). Node dynamics are governed by seven state variables, five of which are local neural-mass states: the mean membrane potential *V*, the mean firing rate *r*, an adaptation variable *u*, and the AMPA and GABA synaptic activations *Sa* and *Sg*. These states are coupled to the two remaining dopaminergic states, extracellular dopamine concentration and D1-mediated neuromodulatory drive on AMPA conductance (Methods, Section 1.3.2) (Fig. 2*a*). Each node accordingly receives afferent drive along three distinct subsystems – excitatory (AMPA), inhibitory (GABA), and pivotally dopaminergic – the last of which does not contribute a parallel current, but rather modulates the AMPA current itself, in that D1-receptor occupancy multiplicatively scales AMPA conductance and thereby renders dopamine a gain-control channel acting upon glutamatergic excitation (Preti et al. 2026; Depannemaecker et al. 2026).

**Fig. 2:**
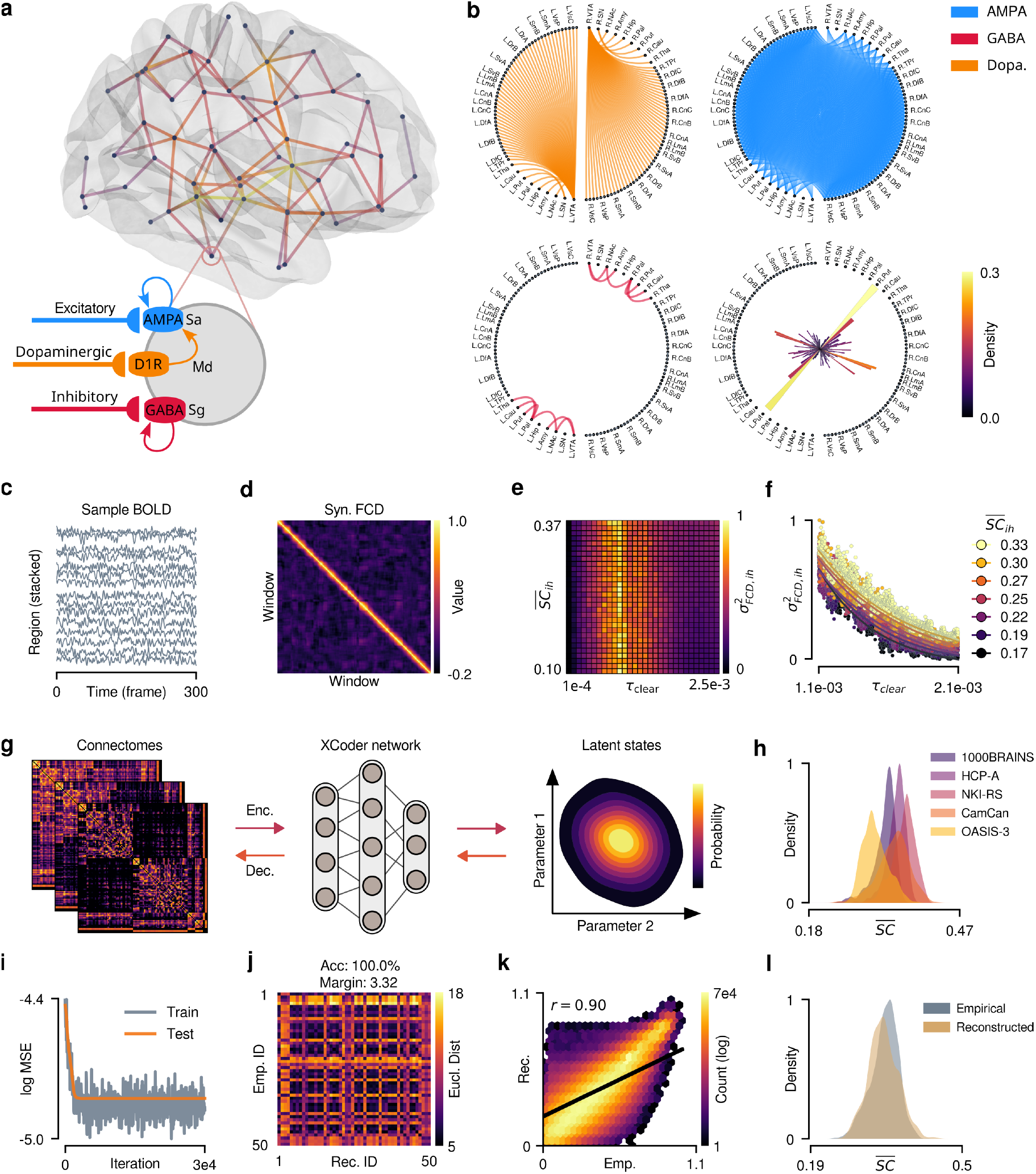
Generative processes making up ADAM. **a**, Schematic of the whole-brain generative model, comprising the AMPA and GABA synaptic conductances and the D1-receptor-mediated neuromodulation of AMPA conductance according to Eq. 1 (full system in Methods, Eqs. 8–9). **b**, Model connectivity masks for dopaminergic (DA), excitatory/glutamatergic (Glu), and inhibitory (GABA) projections, alongside the corresponding regional D1-receptor density heterogeneity vector *Rd*, plotted across the 118-node parcellation. Regional labels are shared across masks and the heterogeneity vector: nodes 1–100 correspond to the Schaefer 17-networks parcellation, nodes 101–114 to the Harvard-Oxford subcortical atlas, and nodes 115–118 to the dopaminergic projection sources (bilateral VTA and SNc) (Bjüorklund and Dunnett 2007). **c**, Sample simulated regional BOLD time series from one representative subject (15 nodes, vertically offset/stacked for visibility purposes). **d**, Sample simulated interhemispheric FCD matrix (axes index sliding time-windows). **e**, Twodimensional sensitivity sweep of the interhemispheric fluidity 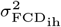 over *τ*_clear_ and mean interhemispheric structural connectivity 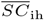, the latter of which was traversed by attenuating only the interhemispheric edges of a single reference connectome across the empirically observed range of 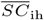 (intrahemispheric and dopaminergic-projection edges left intact; Methods, Section 1.4.1). **f**, One-dimensional sensitivity sweep of 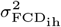 over *τ*_clear_, repeated across 7 1000BRAINS subjects spanning the empirical range of interhemispheric SC and colour-coded by 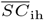 (Methods, Section 1.4.1). **g**, Structural connectivity generative model (CrossCoder network) schematic. **h**, Distributions of harmonized average connectome strength 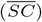 across the five cohorts, illustrating the consistency of cohort-level structural heterogeneity within the shared latent space. **i**, Training and test loss traces of the selected CrossCoder architecture. **j**, Held-out subject-identifiability confusion matrix (*n* = 50 held-out subjects, 1-indexed axes), showing 100% classification accuracy from latent coordinates alone, in conjunction with identifiability margin. **k**, Reconstruction fidelity plot of the selected CrossCoder architecture: log-density hexbin of empirical vs. decoded SC entries, pooled across subjects and non-zero edges (Spearman *ρ* = 0.90). **l**, Distribution comparison of empirical vs. CrossCoder-generated 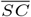.

**Fig. 3:**
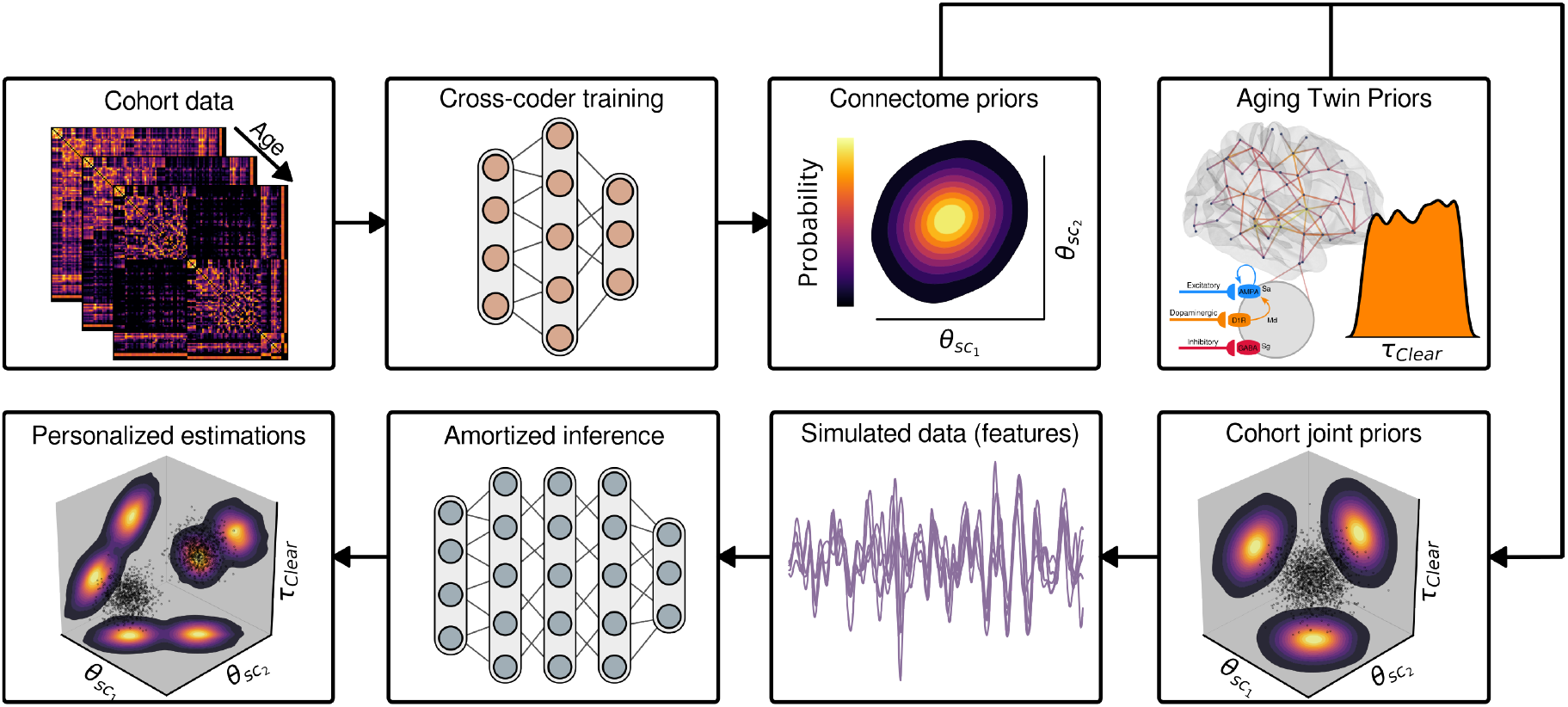
Architecture of ADAM. Empirical structural connectomes from the five aging cohorts are compressed by the CrossCoder into a two-dimensional latent space, yielding a cross-cohort structural prior; this is combined with a normative prior over the dopamine clearance timescales *τ*_clear_, informed by the whole-brain dynamics with underlying dopaminergic synapses (D1-receptor-mediated AMPA modulation and Michaelis–Menten DAT-mediated clearance), to form a joint cohort-amortized prior over the structural and neuromodulatory components. Samples from this prior distribution are decoded into structural connectomes which couple nodes within the aging brain twin to arrive at whole-brain simulations. An amortized conditional density estimation subsequently learns the mapping between multimodal summary statistics from said cohort-scale simulations and model parameters. Applying the trained estimator to each subject’s empirical features in a single forward pass, without any retraining, yields the personalized posterior estimates illustrated in Fig. 4.

Grounded by Michaelis–Menten DAT-mediated reuptake dynamics (Nicholson 1995), the extracellular DA concentration [*Dp*]*e* evolves via:

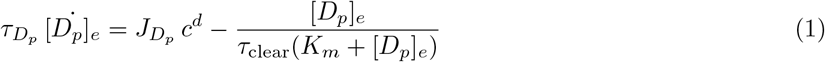

in which *τ*_clear_ – the reciprocal of the maximal DAT reuptake rate, and the only dopaminergic parameter estimated per subject in the sequel – scales the clearance. In this formalism, *τ*_clear_ is not itself a timescale, but the characteristic time constant that sets the timescale over which dopamine is cleared. Larger *τ*_clear_ denotes slower reuptake and, at steady state, elevated extracellular dopamine alongside sustained D1-receptor occupancy, precisely the synaptic environment anticipated for the older brain under preserved synthesis and reduced DAT availability (Karrer et al. 2017). The full system of equations, and the fixed biophysical constants therein, are provided in the Methods (the conventional notation is reparameterized by explicitly representing the dopamine time constant *τD*_*p*_ and substituting *JD*_*p*_ in place of the conventional coupling term *k*). Each subject’s harmonized connectome is partitioned into three neurotransmitter-specific sub-networks supplying the aforementioned afferent drives, by assigning every region a single dominant efferent modality in accordance with established projection neuroanatomy (Preti et al. 2026) (Fig. 2*b*). The glutamatergic sub-network thus spans the bulk of cortico-subcortical edges, the GABAergic sub-network a sparse subset of inhibitory projections, and the dopaminergic sub-network the efferents of four projection-source nodes representing VTA and SNc bilaterally. Connectivity for these four nodes is populated from a tuned template rather than personalized tractography, given the infeasibility of resolving such small and deep nuclei via diffusion imaging at the standard protocols (i.e. 3T) employed for acquisition of all five cohorts in the present investigation. Dopaminergic modulation is further weighted, per node, by the regional D1-receptor density *Rd*, which is derived from an averaged PET receptor-density map (Preti et al. 2026, 2025).

We computed interhemispheric fluidity through simulations obtained from the above model over a grid of *τ*_clear_ and mean interhemispheric structural connectivity 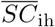 (the latter traversed by attenuating only the interhemispheric edges of a single reference connectome, leaving intrahemispheric and dopaminergic-projection edges intact), so as to obtain an approximate but informative baseline of the model’s response to perturbations in structural connectomes and clearance timescales (Fig. 2*c*–*e*). We obtained a similar non-amortized baseline of interhemispheric fluidity by using each subject’s connectome to conduct individualized simulations (Fig. 2*f*), which will subsequently be used to compare against interhemispheric fluidity estimations of amortized simulations. Simulating over a large range of clearance timescales identified that interhemispheric fluidity varies non-monotonically with *τ*_clear_, following an inverted-U curve (Fig. 2*e*). We thus confined our prior to the falling branch (*τ*_clear_ *∈* [1.1, 2.0] *×* 10^*−*3^) which is aligned with the joint age-related decline of DAT availability and fluidity (Fig. 2*f*). With regards to the second component of the amortized pipeline, we trained a cross-cohort autoencoder (CrossCoder) on empirical structural connectomes harmonized across all five aging cohorts (Methods, Section 1.1.4), so as to learn a two-dimensional shared latent embedding ***θ***SC = (*θ*_SC1_, *θ*_SC2_) (Fig. 2*g, h*).

This autoencoder (due to the existence of a unified parcellation) was trained to minimize reconstruction error while maintaining a well-structured –multivariate normal– latent distribution for employment as a structural prior in the subsequent amortized training (Esmaeili et al. 2026). Our training achieved a reconstruction fidelity of *ρ* = 0.90 (Spearman, over all aggregated non-zero edges, observed versus reconstructed), alongside a 100% accuracy in a held-out subject-identifiability test. The identifiability margin was substantial (avg. off-diagonal minus avg. on-diagonal reconstruction distance 3.32, *≈* 47% larger than the average self-reconstruction distance, Fig. 2*j*), and averages of reconstructed and empirical connectomes were robustly aligned (Fig. 2*k*–*l*). Such robustness is a prerequisite of cohort-scale amortized inference, since non-identifiability of subjects’ latent states equates to a degenerate mapping between macroscale observables and latent state parameters, and with it unreliable personalized estimations. Taken together, our findings corroborate that the CrossCoder training can collapse connectomes into a tractable latent topology while preserving their inter-individual variability.

We combined the aforementioned structural priors ***θ***SC with a uniform prior over *τ*_clear_ to arrive at a (cross)cohort-scale joint prior, formally defined as a three-dimensional parameter space comprising {*θ*_SC1_, *θ*_SC2_, *τ*_clear_}. Next, we sampled from this joint prior, simulated the whole-brain model (*N*sim = 20,000), and finally trained a deep conditional density estimator on the resulting pairs, following our previous protocol of cohort-amortized personalization (Esmaeili et al. 2026; Durkan et al. 2019; Greenberg et al. 2019). Density estimators learn an invertible mapping between model parameters and data features, such that post-training evaluation of empirical data yields posterior estimates in a single forward pass and in a matter of seconds, while remaining ready to draw samples for each new observation (Fig. 3).

### Cohort-amortized personalization of DA clearance timescales

Due to the strong association between 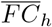 and 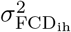 (*r* = 0.68), and our primary interest in dynamic (FCD-derived) rather than static functional summary statistics as a time-varying signature of age-related decline, we opted not to retain 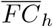 among our minimal set of training features. Our principal criterion for feature selection revolved around (latent) identifiability, where we opted for an informative low-dimensional feature vector (as opposed to a high-dimensional and highly collinear one) to jointly recover the three latent parameters {*θ*_SC1_, *θ*_SC2_, *τ*_clear_} (Section 1.4.1). We therefore chose BOLD-signal standard deviation *σ*BOLD as the complementary dynamic-functional summary alongside 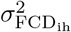, 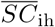, and the signed intrahemispheric SC difference ΔSC, intra (mean left- minus right-hemisphere intrahemispheric SC, homotopically node-matched; see Methods), selected chiefly owing to their capacity in rendering the joint parameter vector identifiable from summary statistics alone. Association of 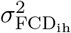, *σ*BOLD, and 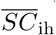 with age (Supplementary Fig. S4) is a welcome, but auxiliary,property of our identifiability-driven selection rather than its rationale. ΔSC,intra best exemplifies this principle, owing its employment to its high identifiability of *θ*_SC2_ despite being a weak and cohort-inconsistent age correlate (see Supplementary Fig. S4).

We next fed the empirical summary statistics of all *N* = 3,650 participants to the trained neural density estimator in order to attain personalized posterior estimations of {*θ*_SC1_, *θ*_SC2_, *τ*_clear_} (Fig. 4). We endeavored to have at least one highly informative data feature for each of the latent parameters, which include 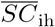, ΔSC,intra, and 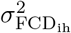 (Fig. 4*a*, see Supplementary Fig. S4 for a thorough visualization of feature-parameter mappings). We report the age dependence of the estimated *τ*_clear_ below, and assess whether its association with cognitive performance is moderated by structural integrity. Given a rigorous training, CrossCoder latents tend to encode explainable and possibly orthogonal summary characteristics of structural connectomes (such as overall SC magnitude, as demonstrated directly below) (Esmaeili et al. 2026).

**Fig. 4:**
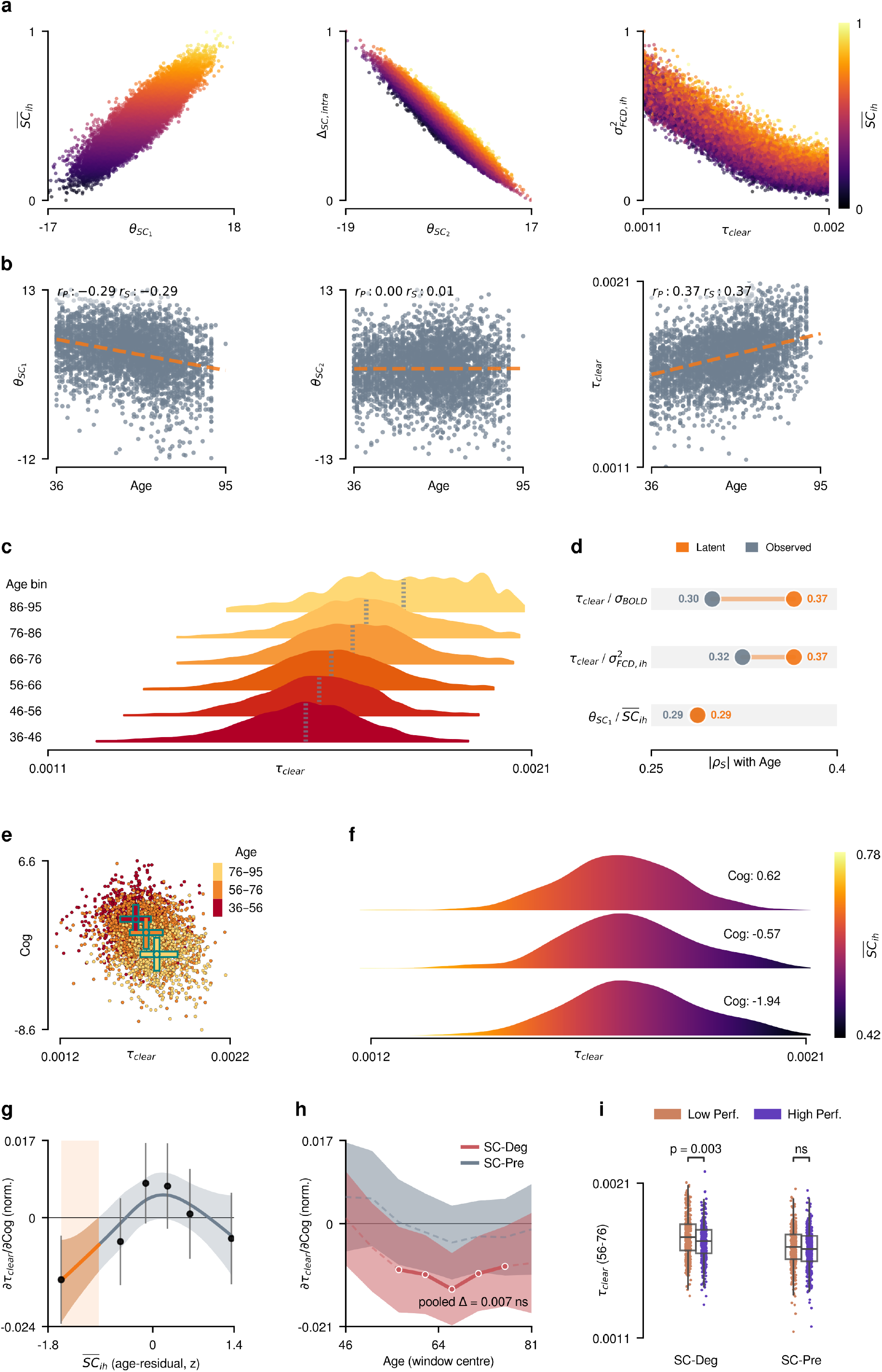
Cohort-amortized personalization of *τ*_clear_ and its relationship to aging and cognition. **a**, Scatter plots of three representative summary statistics against their principal latent counterparts, across the *N*_sim_ = 20,000 training simulations (all colour-coded by 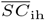): mean interhemispheric SC 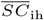 vs. *θ*_SC1_; signed intrahemispheric SC difference ΔSC,intra vs. *θ*_SC2_; and interhemispheric fluidity *σ*^2^ vs. *τ*_clear_. **b**, Correlation of the three inferred posterior means with age: *θ*_SC1_ (*r* = *−*0.29, *p* = 8.4 *×* 10*−*73), *θ*_SC2_ (*r* = 0.00, *p* = 0.81), and *τ*_clear_ (*r* = 0.37, *p* = 6.6 *×* 10^*−*120^). **c**, Age-stratified posterior ridgeline densities of *τ*_clear_ across successive decades from 36 to 95 years. **d**, Comparison of latent (orange) and observed (grey) absolute Spearman correlations with age (|*ρS* |) for three predictor pairs: *τ*_clear_ vs. the BOLD-signal standard deviation *σ*BOLD (latent 0.37 vs. observed 0.30), *τ*_clear_ vs. 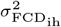 (latent 0.37 vs. observed 0.32), and *θ*_SC1_ vs. *SC*ih (latent 0.29 vs. observed 0.29). **e**, Inferred *τ*_clear_ vs. the global cognitive composite Cog (the first principal component of cognitive performance), stratified into three roughly two-decade-wide age bins (36–56, 56–76, 76–95 years), with per-bin two-dimensional medians overlaid, points colour-coded by age on a colour scale distinct from panel **f. f**, Ridgeline densities of *τ*_clear_ posteriors binned across the range of Cog and colour-coded by 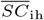. **g**, Model-free shape diagnostic for the cognition*×*structure moderation of *τ*_clear_ (*τ*_clear_ min– max normalized to [0, 1] for panels **g**–**h**, distinct from its raw scale in **b**,**c**,**e**,**f**): local *τ*_clear_ slope on Cog_resid_ estimated within sextiles of age-residualized *SC*ih (points, with standard errors) and smoothed with a penalized natural-cubic spline (curve, shaded bootstrap 95% CI); the orange band marks the region over which the slope’s CI excludes zero. Here Cog_resid_ and SCresid denote Cog and *SC*ih residualized on a quadratic age polynomial and z-scored; the panel axes abbreviate them as *Cog* and *SCih (age-residual, z)*, and their local slope as *∂τ*clear*/∂*Cog. **h**, Sliding-window conditional-slope dissociation: the *τ*_clear_-on-Cog_resid_ slope (with 95% CI) estimated within a 20-year window slid across age, separately for the structurally degraded (SC-Deg) and structurally preserved (SC-Pre) halves of each window; markers denote windows whose CI excludes zero. **i**, Raincloud plots of *τ*_clear_ comparing high and low cognitive performers within the structurally degraded (SC-Deg) and structurally preserved (SC-Pre) groups, restricted to the 56–76 year age band (the range of window centres over which the SC-Deg slope in **h** is significant); *p*-values are from a permutation-based, family-wise-error-controlling (max-*T*) test on the underlying Brunner–Munzel statistic (Methods).

Reliable cohort-amortization therefore necessitates identification of data features which can faithfully embody connectome embeddings. For our purposes, these features consisted of 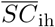 and ΔSC,intra (signed intrahemispheric SC difference) which bore strong associations with *θ*_SC1_ (*r* = 0.89), and *θ*_SC2_(*r* = *−*0.95), respectively, as traced across the training simulations (Fig. 4*a*, left and middle panels; all *p <* 10^*−*50^). The two structural features nevertheless deviate in their age sensitivity, in that a robust age-dependent decline was found in 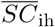 across all cohorts, whereas ΔSC,intra did not exhibit any reliable age-related trend, particularly in cohorts with previously processed data (i.e. 1000BRAINS & HCP-A, Supplementary Fig. S4).

Variations in synthetic 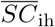 (which has been established as a reliable tracker of age-related decline) also do not seem to have an effect on the value of ΔSC,intra (Fig. 4*a*). Estimation of latent posteriors hence demonstrated a salient decline of the primary structural parameter 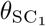 with age (*r* = *−*0.29, *p* = 8.4 *×* 10^*−*^; Fig. 4*b*), and no age-related association in terms of 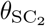 (*r* = 0.00, *p* = 0.81; Fig. 4*b*). Pivotally, our estimation revealed a prominent elevation of the dopamine clearance timescale *τ*_clear_ with advancing age (*r* = 0.37, *p* = 6.6 *×* 10^*−*120^; Fig. 4*b, c*).

The estimated age-related decline of *θ*_SC1_ is concordant with the aforementioned degradation of the structural connectome (and summary statistics thereof) (Fig. 1*h*). This concordance suggests a successful mapping of connectome-oriented decay onto the structural component of the joint parameter space (|*ρS*| = 0.29 obs. vs. 0.29 pred., Fig. 4*d*), rather than a mere assimilation thereof upon the neuromodulatory component (i.e. *τ*_clear_). Our findings with regards to *θ*_SC2_ should in contrast be taken as age-invariance of cross-cohort inter-individual variability, rather than weaker identifiability (Supplementary Fig. S1). Our inferred clearance timescales tracked age more closely than either of the dynamical observables informing them (|*ρS*| = 0.37 for *τ*_clear_, against 0.30 for *σ*BOLD and 0.32 for 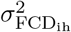 Fig. 4*d*). Taken together, our findings suggest that cohort-amortized partitioning of multimodal observations of the aging brain (into structural and neuromodulatory components) compresses age-relevant information more efficiently than any single observable measure, whilst corroborating previously established DAT attenuation with advancing age (Karrer et al. 2017).

But how do changes in *τ*_clear_ relate to age-related decline of cognitive performance? Younger age bins in general exhibit higher global cognitive composite scores (Cog) and diminished *τ*_clear_ values, with subjects’ estimated clearance timescales within the same cognitive performance bin still displaying a clear association with connectome strength (Fig. 4*e, f*). We thus attempted to isolate the mechanistic influence of structural degradation from that of clearance timescale elongation. Formally, we residualized 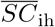 and Cog on a quadratic function of age and fed the resulting Cogresid/SCresid terms into a continuous moderation model, 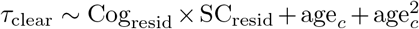 (Methods, Section 1.5). The interaction coefficient in this model directly assesses whether the performance–clearance association hinges on structural integrity. We fit the model over the full age range (36–95 years, *n* = 3,106), which yielded a significant interaction coefficient 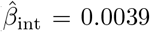. This finding corroborated the dependence of performance–clearance associations on structural integrity (along the direction established by the model-free analysis below). We min–max normalized *τ*_clear_ for this analysis specifically ([0, 1]; distinct from its raw model-calibrated scale elsewhere, e.g. Fig. 4*b, c, e, f*) and z-scored Cog_resid_/SCresid, such that the slope/*β* values are intelligible as a fraction of *τ*_clear_’s range per SD of residualized cognitive performance (Methods, Section 1.5).

First, we implemented a model-free shape characterization of said interaction by binning age-residualized 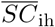 into sextiles, recovering each bin’s local *τ*_clear_ slope on Cog_resid_, and subsequently smoothing across bins through a penalized natural-cubic spline (see Methods). Results of this model revealed the local *τ*_clear_ slope to be significantly negative (bootstrap CI excluding zero) only for the most deteriorated SCs, and inseparable from zero as 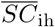 is preserved, thereby suggesting the existence of a degradation-gating mechanism (Fig. 4*g*). In order to investigate the exact age window in which structural deterioration attains significance, we slid a 20-year window over the moderation model. The SC-preserved partition of the cohort exhibited no coupling at any age window (one aggregated/pooled fit over the full group: *β* = 0.0006, *p* = 0.83). The SC-degraded partition, by contrast, bore a run of windows centred between 56 and 76 years during which cognition–*τ*_clear_ coupling was significant (the same pooled fit: *n* = 1,553, *β* = *−*0.0068, *p* = 0.014, corroborated via cluster-based permutation test *p* = 4.6 *×* 10^*−*3^).

We conducted a parallel cluster-based permutation test on the SC-Deg versus SC-Pre slope difference, which did not reach significance in either form (*n* = 3,106; pooled difference = 0.0074, *p* = 0.066; no suprathreshold cluster formed). Additionally, we tested the influence of these analytical choices on this null across 40 combinations of window width (10–30 years), step size (2–5 years), estimator (Huber/OLS) and cluster-forming threshold (|*t*| *>* 1.64 or 1.96), with the pooled difference remaining non-significant throughout (*p* = 0.066–0.074) and no configuration yielding a significant cluster (minimum *p* = 0.098). SC-Deg coupling on the other hand, remained significant in every configuration (cluster *p ≤* 0.023). This may indicate that despite the robustness of SC-Deg coupling itself, the formal between-partition contrast remains underpowered at the present sample size. Coupling was not detected in the oldest (centered 81 years, *p* = 0.124) and the youngest (centered 46 and 51 years, *p* = 0.82 and *p* = 0.29) windows. Informed by the above continuous findings, we conducted an age-constricted stratification (56–76 years), and compared the estimated clearance timescales of high versus low performers by dividing subjects into structurally preserved (SC-Pre) and structurally degraded (SC-Deg) groups via mediansplit of 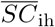. High cognitive performers exhibited significantly lower clearance timescales than low performers in the SC-Deg group (*n* = 795; Brunner–Munzel, *p* = 1.1 *×* 10^*−*3^, Bonferroni-corrected *p* = 2.1 *×* 10^*−*3^, permutation max-*T p* = 2.9 *×* 10^*−*3^, rank-biserial *r*rb = *−*0.13), whereas the same comparison in the SC-Pre group was not significant (*n* = 795, *p* = 0.16, Bonferroni-corrected *p* = 0.32, max-*T p* = 0.29, *r*rb = *−*0.06; Fig. 4*i*). We therefore present the continuous model as our primary estimate of this interaction, and the dichotomized comparison as an auxiliary illustration of it.

Nevertheless, the above findings imply the suitability of *τ*_clear_ as a proxy for quantification of cognitive performance, yet within a specific period of life, and only in individuals with degraded structural connectomes. Our quantification of this decomposition illustrated that across the full cohort, structural deterioration 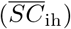 explains a modest but highly significant increment of Cog variance beyond age alone (incremental *R*^2^ = 0.010, *F* (1, 3102) = 45.0, *p* = 2.4 *×* 10^*−*11^). *τ*_clear_ on the other hand, entered as an unconditional (main-effect) covariate on top of SC, itself adding negligible further variance (partial *R*^2^ = 0.00002, *p* = 0.79) which corroborates the abovementioned moderation findings. *τ*_clear_’s pertinence to cognitive performance should therefore not be construed as a global main effect but as a structure-gated and age-bounded interaction; an effect which is only recovered by conditioning in those with considerable SC degradation, and only in a particular age range (windows centred on 56–76 years; Fig. 4*g*–*i*, Supplementary Section S8.).

## Discussion

Age-oriented cognitive decline has been surveyed through myriad hypotheses, and across disparate spatiotemporal brain scales. Nonetheless, multi-scale frameworks attempting to harmonize said accounts have been insufficiently explored. Some neuromodulatory frameworks of aging denote reductions in dopaminergic receptors and transporters, alongside a relatively-preserved synthesis rate in older adults (Karrer et al. 2017). Macroscale inquiries, on the other hand, have alluded to the attrition of interhemispheric structural connectivity and overintegration of resting-state functional networks across the lifespan (Sullivan et al. 2010; Puxeddu et al. 2020; Stumme et al. 2022; Chan et al. 2014; Battaglia et al. 2020). We hypothesize that these accounts denote interrelated perspectives into our brains’ multiscale and multimodal decline, where clearance-related dopaminergic modulation becomes consequential for macroscale dynamics, and for cognitive performance, under structural attrition. Such a harmonization has remained unaddressed and methodologically fraught, due not only to the absence of large-scale multimodal datasets capable of resolving transporter-level kinetics while capturing wholebrain dynamics simultaneously, but also to the lack of neurobiological plausibility of the latent parameters investigated in previous virtual brain twin implementations (Lavanga et al. 2023).

In this study, we employed cross-cohort amortized personalization of a biologically grounded whole-brain model (Gast et al. 2024; Preti et al. 2026; Depannemaecker et al. 2026) to investigate the association between macroscale perturbations in structural/functional connectomes and variations in DA clearance timescales. A population-level range of empirically informed DA clearance timescales was therefore coupled to cross-cohort structural coordinate ranges, to arrive at our amortized distillation of multimodal normative priors. We subsequently selected four macroscale summary statistics, namely interhemispheric fluidity 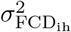, BOLD-signal standard deviation *σ*BOLD, mean interhemispheric SC 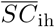, and the signed intrahemispheric SC difference ΔSC,intra. These four data features jointly identify the neuromodulatory and structural parameters of a single amortized estimator, bypassing the need for per-subject and per-cohort retraining. Recapitulating the population-level decline of DATs reported empirically (Karrer et al. 2017), personalized dopamine clearance timescales *τ*_clear_ recovered through this approach exhibited a prominent elevation with advancing age, with this elevation exceeding in magnitude the age association of any single observable data feature which informed the aforementioned probabilistic inference. Only the target neuromodulatory parameter exhibited an advantage over its corresponding observables, and the principal structural coordinate *θ*_SC1_ remained essentially on par with its observable counterpart.

These recovered timescales illustrated a context-oriented disposition, in that elevated *τ*_clear_ separated low from high performers specifically in structurally degraded older adults across a bounded window spanning the 56– 76 year band, while leaving structurally preserved individuals of matched age undifferentiated. In concert, these findings recast a parameter that prior frameworks could only approach phenomenologically as a mechanistically interpretable, scalable, and transferable readout of dopaminergic aging.

### Biologically plausible partitioning of a phenomenological proxy

Switching from connectome coupling scalars in prior frameworks to the current *τ*_clear_, as a potential contributor to age-related decline, does not constitute a mere re-parametrization. Scalars remain phenomenological by definition, and their explanatory value is thus limited, which amounts to a dearth of explicit biological signaling pathways, much less a dopaminergic one. In contrast, our estimation of dopamine clearance timescale is informed by previously established population-level quantification of age-related DAT availability (Karrer et al. 2017) and is anchored in a mechanistic representation of structural decay. The inferential robustness of our latent-over-observed approach stems from this very biologically plausible basis. The CrossCoder coordinates revolve around structural summary statistics, whilst clearance timescales are underpinned by functional ones. The two therefore remain well-partitioned within the personalized estimates, as opposed to collapsing on a singular parameter. The amortized estimation outlined here, to the best of our knowledge, constitutes the first decomposition of accessible macroscale observables in the aging brain into interpretable neuromodulatory and structural components. Cohort-amortization is thus indispensable to the present investigation for a variety of reasons: previous efforts were conducted in a non-amortized manner, which necessitates a computationally expensive re-execution of per-subject simulations and parameter optimization. Hence, the compute budget in this case linearly scales with the number of subjects (such as extension to multiple lifespan cohorts). Conversely, cohort-amortized formalism enables one to train a shared estimator on a single set of cross-cohort simulations, and to obtain accurate personalized estimations on the scale of seconds at inference time (Esmaeili et al. 2026). Cohort-amortization’s primary endowment is more substantive than mere alleviation of computational cost, since a unified estimator at the scale of cohorts is the most principled conduit for preserving similarly cohort-quantified (and not individually measurable) DA clearance timescales, whose normative range is itself inherited from cohort-pooled DAT availability (Karrer et al. 2017). Moreover, this approach yields personalized estimates conditioned on subjects’ data features through a unified cohort-scale estimator.

This amortized and practical framework was initially inspired by our own experience of inter-institutional datasharing boundaries throughout this endeavor, which, we believe, heavily constrains the translational impact of computational forays into the aging brain. Interoperability, multimodality, and atlas-agnosticism of the general cohort-amortization schema empower lifespan researchers to bypass exchange of raw data, and hurdles thereof, and instead distribute their trained estimators across sites. ADAM also allows for the conduct of synthetic neuromodulatory experiments, generation of synthetic aging cohorts, and efficient benchmarking of disparate macroscale/cognitive markers against a standard mechanistic data generative process.

### Moderation of dopaminergic clearance through structural integrity

As established in Fig. 4*g*–*i*, both the continuous and stratified quantifications of *τ*_clear_ converge to a shared subset of profiles and stages for which dopaminergic clearance could be consequential in the partitioning of cognitive decline; within the 56–76-year age band, elevated *τ*_clear_ marks the lower-performing stratum principally among individuals with prominent structural degradation. Structural degradation should therefore be viewed as a permissive context that unearths such a neuromodulatory disparity, rather than the definitive cause of it. One promising rendition of said disparity is seen through the lens of neuronal gain control (Li et al. 2001; Li and Rieckmann 2014). Dopaminergic regulation under this view modulates the gain of neuronal stimulus-response mappings, resulting in changes to neural representation diversity and fidelity (i.e. signal-to-noise ratio) of cortical dynamics. For the present account, the level of that gain is set by prevailing receptor occupancy, which the balance between dopamine release and clearance itself governs (Dreher and Burnod 2002). Whether elevated dopaminergic input enhances or degrades that ratio primarily depends on the baseline levels of dopamine, as opposed to the increment itself (Cools and D’Esposito 2011; Kroener et al. 2009). Indeed, older adults with enhanced retention of striatal D1R availability display both better working memory and preserved functional segregation (Pedersen et al. 2023). Furthermore, diminished prefrontal and caudate D1R availability has been purported to signal poorer memory and impoverished neural gain control (Korkki et al. 2025).

While these findings establish D1-mediated gain control as a promising contributor to age-related decline, they exclude the concurrent reduction in DATs and the attenuation of interhemispheric connectivity, and are bounded by receptor availability as its sole synaptic underpinning. ADAM’s formulation, on the other hand, augments the previous accounts of dopaminergic decline through a biologically plausible representation of clearance *τ*_clear_ (depicted in Eq. 1), and operates within the same framework through an axis distinct from structural degradation. In this formulation, clearance-dependent D1 modulation of AMPA conductance influences the excitatory drive of each neural population. Within the confines of our model configuration, longer DA clearance timescales therefore elevate D1-modulated AMPA drive and in turn, excitability. Elevated excitability allows for better regional coupling via structural connectivity. On a diffuse spectrum of reuptake timescales, this enhancement of structure-function coupling carries time-resolved synchronization to an initial peak. Populations beyond this peak exhibit progressively diminished alternation between high- and low-activity states (Hansen et al. 2015), and hence reduced functional reconfiguration (Supplementary Section S4.). Attenuated structure-function coupling present under interhemispheric attrition (Zhang et al. 2024) hence allows for differences in dopaminergic modulation to be cognitively consequential.

Posterior stratification of clearance timescales presented here may underlie inter-individual heterogeneity in the dynamic reconfiguration of functional connectivity. Viewed in this light, *τ*_clear_ represents a candidate through which the gain-control theory elucidates cognitive performance disparity, and provides one with a biologically plausible mechanism that molecular imaging by itself could neither localize nor measure across large-scale cohorts. The present work therefore does not portray a main effect of *τ*_clear_ upon cognition, but rather an *interaction* with structural integrity. Indeed, structural attrition has been shown to track cognitive decline longitudinally within individuals (Pedersen et al. 2021), an attrition which reduces network dynamical reserves, thereby permitting residual dopaminergic modulation to exert a greater influence on neural dynamics.

Our continuous moderation model estimates this influence directly, as a single coefficient, while the model-free shape check confirms that the slope meaningfully diverges from zero only at the most degraded connectomes. That structure-dependence is itself age-constrained, emerging in midlife and weakening past the mid-70s (where it favours neither performance stratum). A late-life convergence of dopaminergic and structural decay may therefore serve as a reasonable account for this trend, and is compatible with previously reported attenuations in DAT (Karrer et al. 2017). Put together, the amortization afforded by ADAM reconciles the accounts of agerelated dopaminergic (Pedersen et al. 2023; Korkki et al. 2025) and structural (Pedersen et al. 2021) decline in a unified mechanistic framework, and outlines structural deterioration as the manifold upon which DA clearance timescales may be cognitively consequential.

## Limitations and scope

The scope of the work outlined above is demarcated by four constraints. For one, we have chosen to deliberately omit the modeling of D2 signaling pathways in addition to the serotonergic subsystem, which is seen in the full neuromodulatory formulation of the Schizophrenia Virtual Brain Twin (Preti et al. 2026). Such a reduction concentrates inference on the most thoroughly characterized molecular aging signature, and eschews identifiability issues pertaining to probabilistic inference of neuromodulatory factors affecting similar additive currents via whole-brain summary statistics (which are inherently insufficient to disentangle/inform said degeneracy). Yet the serotonergic system, akin to every other biological subsystem, is not exempt from age-related deterioration (Kanel et al. 2023). Augmenting ADAM to jointly estimate 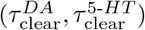 thus constitutes a natural unfolding of the framework which ADAM’s architecture accommodates without structural modification, and is only constrained by the trade-off between mechanistic completeness and posterior identifiability.

The second limitation concerns the mean-field delineation of DAT-mediated clearance processes and a lack of D1R personalization. Inferred *τ*_clear_ constitute effective, spatially aggregated (i.e. mean-field) clearance timescales, but do not resolve the regional (e.g. striatal and prefrontal) assortment documented by PET studies (Karrer et al. 2017). Due to the absence of data required for personalization of receptor densities, our chosen D1-density map *Rd* is likewise fixed to a population average. As such, the age-related changes mentioned above should not be construed as a claim of anatomical uniformity, but rather as a first-order reduction and principled rescaling of clearance across DA-modulated regions/nodes. In the absence of auxiliary constraints such as direct DAT-PET measurements, node-resolved estimation of *τ*_clear_ would escalate the effective dimensionality of the inference beyond the reach of the current observation model, and would –similar to the previous limitation– likely result in non-identifiability at the subject scale. This line of inquiry should naturally be extended by a PET-informed validation of personalized *τ*_clear_ estimates against measured DAT availability and receptor densities in the same subjects.

The third issue stems from the inclusion of cross-sectional rather than longitudinal data across all five cohorts. All age-dependent declines observed via the employment of such datasets ought to be portrayed as age-*associated* rather than age-*caused* effects, so as to not conflate cohort-specific recruitment biases with genuine aging dynamics. And while this is an extensive issue in aging neuroimaging datasets at large, it bears particular pertinence for investigations whose central claims hinge on relatively restricted transition windows, such as the current study. ADAM heavily reduces per-subject inference cost, which allows for repeated-measures designs at the scale of cohorts and populations. Longitudinal renditions of cohort-amortized personalization could therefore rectify this ambiguity directly, and would serve as a natural extension of our cross-sectional approach.

Finally, we should note that our approach posits the dopaminergic clearance timescale not as the sole contributor to the age-related decline of dynamical reserves considered here, but as an identifiable and promising candidate. Age-oriented deterioration in neural systems has been hypothesized to stem from myriad mechanistic sources, such as elevated aperiodic (broadband) neural noise, and reduced cortical GABA concentration (Lalwani et al. 2019), mechanisms which partially overlap with, but ought not to be conflated with, the dopaminergic gain-control account exemplified here. In addition, the link between *τ*_clear_ and macroscale decline is causal only *with respect* to the outlined generative processes. Simply put, the estimated *τ*_clear_ can only go so far as to substantiate the consistency and mechanistic adequacy of the macroscale summary statistics (Stumme et al. 2022). Therefore, the approach envisioned by our efforts here can be best described as a framework which –in the absence of large-scale in-vivo observations of DA clearance– presents researchers with an experimentally falsifiable and mechanistically explicit in-silico hypothesis.

Despite the above constraints, the endeavor introduced here demonstrates that ADAM’s biologically plausible, cohort-amortized personalization can recover subject-specific estimates of an effective dopaminergic clearance timescale from coarse-grained yet multimodal neuroimaging. Clearance timescales recovered through such means trace a general aging trajectory, and a structure- and age-conditional association with cognitive performance, with a fidelity that surpasses the resolving power of any single observable data feature considered here. This mechanistic resolution allows for efficient and scalable disentanglement of high and low cognitive performers across disparate aging cohorts, against a common structural background.

## 1 Methods

### 1.1 Data cohorts

Empirical macroscale observables employed in this study are computed from five independent aging cohorts, comprising the first visits of each participant: 1000BRAINS; HCP-Aging (HCP-A); the NKI-Rockland Sample (NKI-RS); Cam-CAN; and OASIS-3. Following quality control and the matching of subjects possessing usable diffusion-weighted, resting-state functional, and demographic data (Section 1.1.4), we retained samples of *N*1000BRAINS = 841, *N*HCP-A = 1,208, *N*NKI-RS = 544, *N*Cam-CAN = 511, and *N*OASIS-3 = 546, yielding a combined *N* = 3,650 participants spanning an age range of 36–95 years.

#### 1.1.1 1000BRAINS

Grounded in the Heinz Nixdorf Recall Study of the University of Duisburg-Essen (Germany), the 1000BRAINS project is a population-based inquiry into healthy aging, aiming to characterize the inherent variability of neural structure and function relative to behavioral, environmental, and genetic determinants (Caspers et al. 2014). We refer to the original cohort description (Caspers et al. 2014) for full acquisition specifics (3T Siemens Tim-TRIO; 32-channel head coil) and the established structural, diffusion, and resting-state protocols. This cohort forms the basis of our initial investigation, due to the availability of parcellated BOLD and structural data for our desired hybrid atlas. From the latest iteration of the 1000BRAINS dataset accessible to us, we extracted solely the first visit for participants exceeding 36 years of age for whom we confirmed the availability of high-quality diffusion and resting-state fMRI. Diffusion processing was conducted using a multi-shell HARDI approach, implementing multi-tissue constrained spherical deconvolution and iFOD2 probabilistic tractography within MRtrix3 (incorporating SIFT2 streamline weighting) to construct connectivity matrices over the 100-region/17-network Schaefer parcellation (Schaefer et al. 2018), supplemented by the 14 Harvard-Oxford subcortical regions (resting-state time series were available for the 100 cortical parcels only), following the acquisition and processing conventions described in Caspers et al. (2014). Resting-state fMRI data (gradient-echo EPI; TR = 2200 ms; 300 volumes; *≈*11 min) were processed using fMRIPrep (Esteban et al. 2019) and subsequently denoised via the Nilearn interface against fMRIPrep confounds. In terms of data variants, we selected the preprocessing variant which corresponds to Nilearn’s *simple* confound-removal strategy (Abraham et al. 2014), for the BOLD time-series, which encompasses cosine-basis high-pass filtering, 24-parameter motion regression (Friston et al.

1996), a 0.1 Hz low-pass filter, and mean WM/CSF regression (excluding global signal regression and scrubbing; V4 variant), having discarded the first four volumes of each run (296 retained), while opting for the raw SIFT2-weighted connectomes as the primary SC matrices for the connectome harmonization purposes.

#### 1.1.2 HCP-Aging (HCP-A)

The Lifespan Human Connectome Project in Aging (HCP-A) recruits community-dwelling adults across the lifespan, employing acquisition and processing protocols shared with the broader Lifespan HCP initiative (Bookheimer et al. 2019; Harms et al. 2018). HCP-A constituted the earliest of our four in-house-analyzed cohorts, with its minimal-preprocessing release helpfully already harboring eddy- and susceptibility-corrected diffusion data (with fiber-orientation distributions fit via FSL (Jenkinson et al. 2012) `bedpostX`) and restingstate fMRI resampled into CIFTI grayordinate (`dtseries`) space. We leveraged these data to bypass the majority of the steps involved in our in-house tractography and parcellation pipeline relative to the unified approach taken with NKI-RS, Cam-CAN, and OASIS-3 (Section 1.1.3). To generate structural connectomes, we executed GPU-accelerated probabilistic tractography (`probtrackX2 gpu`, network mode, one-way condition) on the precomputed `bedpostX` fiber-orientation samples. We seeded this process from grey-matter voxels of the 100-region Schaefer-17Networks parcellation (Schaefer et al. 2018), supplemented by 14 Harvard-Oxford subcortical regions to maintain atlas identity across cohorts prior to CrossCoder harmonization. We set the tracking parameters to a curvature threshold *c* = 0.6, 5,000 seed samples per voxel (*−P*), 3,000 propagation steps (*−S*), and a step length of 0.5 mm. We chose this step length in light of our per-cohort tuning with respect to cohort-specific number of diffusion-encoding gradient directions for each acquisition, which harmonizes the resulting connectome density across cohorts. Parcellated BOLD time-series were derived directly in grayordinate space owing to the availability of the resting-state data as ICA-FIX-denoised (`clean`), MSMAll-registered `dtseries` CIFTI files. In contrast with the volumetric registration and extraction methods we employed for the other three cohorts, we extracted per-node time series by parcellating each run with Connectome Workbench’s (Marcus et al. 2011) `wb command -cifti-parcellate` onto the same Schaefer-17Networks atlas (resampled to grayordinate `dlabel` space). Finally, we concatenated the runs across available phase-encoding orientations.

#### 1.1.3 NKI-Rockland Sample (NKI-RS), Cam-CAN, and OASIS-3

The remaining three cohorts comprise NKI-RS, Cam-CAN, and OASIS-3, the demographics and acquisition descriptions for which can be found in their respective original publications (Nooner et al. 2012; Shafto et al. 2014; Taylor et al. 2017; LaMontagne et al. 2019). We employed a unified preprocessing and parcellation of diffusion-weighted and resting-state functional data of said cohorts, in order to maintain control over our desired parcellation scheme and tractography parameters across otherwise heterogeneous acquisition protocols.

##### Parcellated structural connectivity estimation

As a first step, T1-weighted images were skull-stripped and segmented into grey matter, white matter, and cerebrospinal fluid compartments. We then warped these compartments into MNI152 space alongside the atlas, and brought both into each subject’s native diffusion space by composing that warp with a T1-to-diffusion registration. This registration was boundary-based for Cam-CAN/NKI-RS and a mutual-information-based affine one for OASIS-3 (in order to accommodate the lower number of diffusion-encoding gradient directions for this cohort). Subsequently, we corrected the diffusion data for eddy-current and motion artifacts utilizing FSL `eddy cuda` (Cam-CAN, NKI-RS), while we applied a classic linear correction pipeline for OASIS-3. Due to the heterogeneity of available reversed phase-encoding acquisitions across all three cohorts, we did not apply susceptibility-induced distortion correction with FSL `topup`. Fiber orientation distributions were estimated via FSL `bedpostX gpu` and probabilistic tractography conducted via `probtrackX2 gpu`. These models were seeded from grey-matter voxels of each of the 100 Schaefer-17Networks cortical parcels (Schaefer et al. 2018), supplemented by 14 Harvard-Oxford subcortical regions. We endeavored to retain a unified set of parameters across cohorts, but opted to vary some of them due to their inherent differences in acquisition configurations. All three shared a curvature threshold of *c* = 0.2, 5,000 seed samples per voxel (*−P*) with 3,000 propagation steps per sample (*−S*), modified-Euler streamline integration with enabled loop-checking, and a fibre-volume-fraction threshold of 0.01. Following our rationale for HCP-A (Section 1.1.2), we chose a step length of 1.0 mm for Cam-CAN/NKI-RS and 2.5 mm for OASIS-3. Grey-matter and CSF thresholds (GM = 0.25, CSF = 0.20) were applied and a 20-voxel minimum seed-volume criterion uniformly across the three cohorts to exclude regions compromised by atrophy or partial-volume effects; an atlas-ROI fallback was additionally implemented for nodes failing the minimum-volume criterion under substantial cortical atrophy. Owing to the inherent asymmetry of connectomes obtained from probabilistic algorithms, we symmetrized the streamline counts between each pair of regions and assembled them into a 114 *×* 114 connectivity matrix (comprising 100 cortical and 14 subcortical nodes) for every subject, setting the diagonal to zero and normalizing edge weights by their cohort-wise maximum (previous to cross-cohort harmonization).

##### Resting-state BOLD time-series

Resting-state fMRI data were processed via the same anatomical registration as our structural pipeline, directly reusing its T1-derived transforms as opposed to re-estimating them. Said data were motion-corrected and denoised through cohort-specific framewise-displacement (Power et al. 2012, 2014) scrubbing thresholds (FD = 0.40 mm for Cam-CAN, 0.30 mm for NKI-RS, 0.50 mm for OASIS-3), tolerated scrubbed-volume fraction maxima (0.30 for Cam-CAN, 0.25 for NKI-RS, 0.30 for OASIS-3). We also opted for cohort-specific grey-matter masking thresholds (0.25 for Cam-CAN, 0.30 for NKI-RS, 0.15 for OASIS-3) so as to retain comparable proportions of usable volumes irrespective of the cohorts’ baseline image quality and disparate motion distributions. Confound regression comprised a 24-parameter motion expansion (Friston et al. 1996) alongside the aforementioned spike regressors. We opted for a more stringent set of preprocessing steps for OASIS-3, due to its coarser acquisition, supplementing the above with mean white-matter and cerebrospinal-fluid signals and a zero-phase 0.01–0.1 Hz Butterworth band-pass. Finally, regional mean time-series were extracted over the same 100-region Schaefer parcellation employed for structural connectivity, and any runs containing fewer than 296 usable volumes were dropped.

#### 1.1.4 Cross-cohort feature harmonization

Disparities in acquisition protocols, tractography pipelines, and tractography/streamline-count conventions can result in systemic heterogeneity in raw connectome edge-weight distributions. In that light, we opted not to directly pool raw structural connectomes across cohorts, and instead harmonized these connectomes prior to CrossCoder training. We first log10(1 + *w*)-transformed subjects’ connectomes, and subsequently applied a per-cohort affine transform (a single scalar and offset, shared across all subjects within that cohort) to match the cohort’s log-transformed mean and maximum edge weight to those of the 1000BRAINS reference cohort, consequently aligning connectome magnitudes across cohorts while fully preserving intra-subject and inter-subject edge weight distributions. Subjects whose summary statistics identified them as outliers were then excluded from subsequent analyses. This outlier detection revolved around a median-absolute-deviation criterion and with its tolerance set per-cohort and per-modality (i.e. FC/SC), on account of significant inter-cohort variability in acquisition quality and raw feature distribution dispersion (*n*MAD = 6.0–8.0 for SC-derived features, 5.0– 6.0 for FC-derived features, alongside a cohort-level criterion of *n*MAD = 4–7). Only a select few subjects were excluded per cohort (1000BRAINS: 19/860, 2.2%; HCP-A: 7/1,215, 0.6%; NKI-RS: 11/555, 2.0%; Cam-CAN: 3/514, 0.6%; OASIS-3: 6/552, 1.1%). Such an exclusion criterion not only safeguards agnostically against gross acquisition or processing failures, but also underpins the normalization of empirical summary statistics and their alignment to (similarly normalized) synthetic data features in cross-cohort analyses such as ours. We subsequently employed a two-pass robust-affine harmonization sequence, this time targeting summary-statistics, so as to align feature central tendency and scale to a common reference, with the first pass provisionally harmonizing both connectomes and extracting summary statistics, and flagging the abovementioned outliers, whilst the second re-harmonizes (at both the connectome and feature level) the surviving subjects. We derived the reference location/scale statistics for all data features from the 1000BRAINS cohort alone, with the remaining cohorts being projected onto that reference rather than onto a pooled cross-cohort target. Age-residualization was intentionally omitted to ensure any age-dependent effect observed within the harmonized summary statistics (Fig. 1*h*) represents per-cohort –albeit dimensionless– ground-truths. We concluded the feature computation procedure by conducting min–max normalization of all summary statistics prior to any subsequent analysis or inference (Section 1.4).

#### 1.1.5 Reduced representations of cognitive performance

We were not able to obtain the array of cognitive scores from the NKI-RS cohort by the time of completion of the current investigation, and as such, all measures of cognition (and associations thereof) concern the remaining four cohorts. We explicitly excluded crystallized achievement measures wherever a cohort administered one (the 1000BRAINS vocabulary test and the Cam-CAN proverbs task). The retained batteries nonetheless include measures sensitive to acquired verbal knowledge, such as category fluency and famous-face naming, so that Cog ought to be read as a broad cognitive-performance score rather than a pure fluid-cognition score. In order to eliminate task-specific scaling, we z-scored raw task scores per-cohort (clipping all extreme outliers at *±*4 SD). Each task’s Spearman correlation with age was calculated, with subsequent inversion of the sign of any task correlating positively with age, resulting in a consistent orientation across all tasks and cohorts. Higher values in all tasks would hence indicate enhanced performance following this measure (with most tasks not changing sign due to them measuring accuracy or performance). These directions were determined once and held fixed thereafter. We computed the first principal component of these z-scored, sign-aligned task-level scores to arrive at a unified global cognitive composite Cog following the multivariate reduction strategy of Stumme et al. (2022), and opted to pass over domain-level aggregation for all cohorts. Finally, we flipped the resulting component’s sign if anticorrelated with the mean of the aligned tasks. Thorough account of per-cohort task inventories and validation against raw task-level age-correlations can be found in Supplementary Section S7..

#### 1.1.6 Empirical observations of DAT decline

The empirical data with respect to PET-SPECT measurements of DAT decline were sourced from a subset of the studies analyzed in the meta-analysis of Karrer et al. (2017), and –in addition to receptor-density and synthesis capacity variables– are openly available from the authors’ data repository at osf.io/6uf82. Individual withinstudy measures were z-scored and subsequently pooled to arrive at a close approximation of the cross-study DAT decline seen in the original publication (Fig. 1*b*; Spearman *ρ* = *−*0.63, *n* = 838). It should be noted that this approximation does not aim to replicate the Fisher-z-transformed, weighted random-effects meta-analysis reported by Karrer et al. (2017), which yielded a striatal DAT value of *r* = *−*0.68 (33 studies, *n* = 1,046), as the authors themselves allude to the emergence of such discrepancies between the abovementioned pooling and consolidation of per-study effect sizes. Our motivation for presenting the data visualized in Fig. 1*b* is to solely demonstrate the approximate magnitude and qualitative direction of age-dependent DAT attenuation.

### 1.2 CrossCoder training and validation

Structural connectomes were harmonized across cohorts into unified low-dimensional embeddings comprising ADAM’s structural Bayesian prior. The cohort-amortized personalization framework accommodates training data comprising multiple per-subject atlases or a single one, the latter of which equates the network to an autoencoder. Although the present work imposes a unified parcellation across all five cohorts (Section 1.3.1), we opted to retain the CrossCoder nomenclature for consistency with Esmaeili et al. (2026). Accordingly, each subject’s (118 *×* 118) connectome was flattened into its upper-triangular vector **c** *∈* R^*d*^ (*d* = 118 *·* 117*/*2 = 6,903) and mapped through a deterministic linear autoencoder:

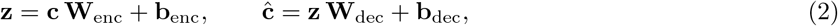

Connectomes were *center* normalized (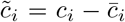, with non-negativity enforced solely during decoding), and fed as input to train the network by minimizing reconstruction MSE (Adam optimizer with learning rate 10^*−*5^, minibatch size 64) over 30,000 iterations. We divided the connectome data array into an 80/20 train/test split, and guaranteed a permuted representation of all cohorts by shuffling the subjects (dimension) prior to this division. We compared candidate latent dimensionalities *n*lat *∈ {*2, 3, 4*}* based on confusion rate, subject identifiability, and reconstruction fidelity (Fig. 2*i*–*k*, Supplementary Fig. S2). The trained CrossCoder –on held-out subjects– achieved a Spearman reconstruction fidelity of *ρ ≥* 0.90 between empirical and reconstructed SC edge weights and 100% subject-identifiability accuracy (each subject’s own reconstruction being its nearest neighbour, in Euclidean edge-weight distance, among the reconstructions of 50 randomly drawn held-out subjects) across all trained dimensions (Supplementary Fig. S2). We thus opted for *n*lat = 2 as the most parsimonious architecture, since amplifying *n*lat beyond latent spaces with two dimensions failed to provide meaningful gains in subject identifiability or reconstruction fidelity. Finally, we fit a multivariate normal distribution *N* (***µ***SC, **Σ**SC) over the held-out (test-split) latent dimensions to attain the structural component of our joint Bayesian prior (Section 1.5).

### 1.3 Dopaminergic whole-brain model

In this section we delineate the forward model at the core of ADAM including regional dynamics, neurotransmitter-specific coupling, and fixed biophysical constants. ADAM refers throughout to the model and inference pipeline developed here, which is built upon the general cohort-amortized formalism of Esmaeili et al. (2026).

#### 1.3.1 Parcellation and connectivity masks

Nodes within the whole-brain model are coupled via a 118 by 118 parcellated connectome comprising the 100-parcels Schaefer-17Networks cortical atlas (Schaefer et al. 2018), 14 subcortical regions from the Harvard-Oxford atlas, and 4 dopaminergic projection nodes (bilateral VTA and SNc). We subsequently partitioned each connectome into three neurotransmitter-specific sub-networks – excitatory glutamatergic (*C*^*e*^), inhibitory GABAergic (*C*^*i*^), and dopaminergic (*C*^*d*^) – following the partitioning schema employed in Preti et al. (2026). Said decomposition was based on established anatomical pathways (Zhang et al. 2017; Lanciego et al. 2012), including: 1) the classical GABAergic circuitry of the basal ganglia, excluding the subthalamic nucleus, which was not segmented, 2) GABAergic projections from the basal ganglia to both the VTA and SNc, 3) Dopaminergic projections from the VTA to the cortex and from the SNc to the basal ganglia. Each region was allotted a single dominant efferent modality grounded on established neuroanatomical projection orientation. In this scheme, only the four bilateral VTA/SNc projection-source nodes are themselves excluded from receiving dopaminergic input (i.e. *Rd* = 0), with the remaining 114 cortical and subcortical nodes all receiving dopaminergic input, albeit at widely disparate levels. We accordingly scaled *Rd* at these 114 nodes based on cohort-level PET receptor-density maps as calculated and presented by Preti et al. (2026) from underlying PET data published in Kaller et al. (2017) (rescaled to [0, 0.3] relative to the chosen baseline of *Bd* = 1), reflecting the previously established strong correspondence between receptor density and dopaminergic structural connectivity reported by Preti et al. (2026) (*≈* 76% shared variance). The tonic excitatory drive *I* in a subset of basal-ganglia targets (bilateral pallidum) was enhanced (*×*1.3 relative to baseline) to restore plausible BOLD dynamics at these nodes: in the absence of this adjustment, GABAergic input reaching these nodes (Zhang et al. 2017; Lanciego et al. 2012) would drive their simulated activity towards quiescence. With regards to personalized probabilistic tractography (Section 1.1.3), we only tracked a 114 *×* 114 structural connectome for each subject (covering only the Schaefer cortical and Harvard-Oxford subcortical nodes). We did not estimate SNc/VTA connectivity due to the deep location, small size, and limited SNR of these nuclei with respect to the standard 3T protocols used for data acquisition across all five cohorts. This is a resolution constraint rather than one stemming from diffusion tractography in general. We instead opted to infill the four DA projection-source edges of the 118 *×* 118 SC from a template connectome which was tuned to the model’s dynamical working regime. This template was scaled to each cohort’s empirical connectome magnitude using the cohort-wise median of per-subject maxima, and integrated around each subject’s unmodified 114 *×* 114 tracked connectome.

#### 1.3.2 Mesoscale local dynamics

We represent mesoscale (i.e. nodal) dynamics via a next-generation neural mass model which originates from a mean-field approximation of a spiking neural network of Izhikevich neurons featuring conductance-based AMPA and GABA synapses (Gast et al. 2024). Each node bears state variables of (*r, v, u, sa, sg*, [*Dp*]*e, MD*1), representing the mean firing rate, mean membrane potential, an adaptation variable, AMPA and GABA synaptic activations, the extracellular dopamine concentration, and the D1-mediated neuromodulatory drive on AMPA conductance, respectively:

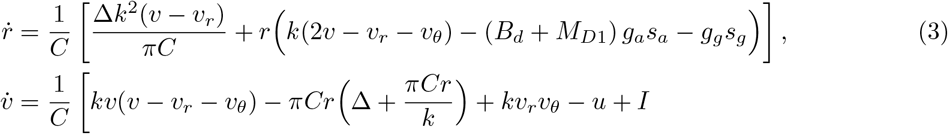

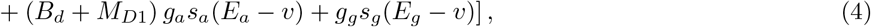

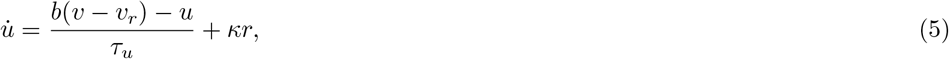

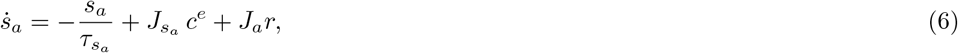

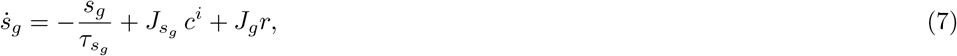

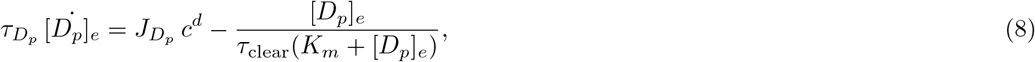

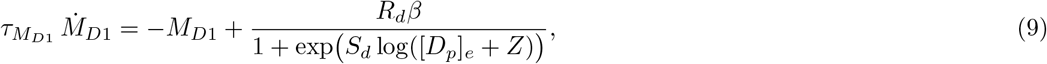

where *C, k*, Δ, *vr, vθ, Bd, ga, gg, Ea, Eg, τu, b, κ, τs*_*a*_, *τs*_*g*_, *Ja, Jg, Js*_*a*_, *Js*_*g*_, *JD*_*p*_, *Km, τD*_*p*_, *τM*_*D*1_, *Sd*, and *Z* constitute fixed biophysical constants (parameterized following Preti et al. (2026); Depannemaecker et al. (2026); Gast et al. (2024) and detailed in Section 1.3.5). Basal AMPA-conductance offset is represented via *Bd* and is tasked with ensuring persistent excitatory drive in the absence of dopaminergic modulation. The parameter *β* scales receptor-sensitivity (which is fixed to 1 in the present work) but in the presence of personalized receptor maps, enables future in-silico inquiries.

Equation (8) delineates the Michaelis–Menten reuptake formalism of Nicholson (1995); Depannemaecker et al. (2026), with our target parameter, the effective clearance timescale *τ*_clear_ inversely scaling the transportermediated reuptake. Elevation in *τ*_clear_ signifies slower DAT-mediated clearance, which in turn enhances [*Dp*]*e* at steady state and increases D1-receptor occupancy and with it *MD*1 via Equation (9), thereby representing the empirical signature of diminished DAT availability despite preserved DA synthesis (Karrer et al. 2017). Due to the stochastic nature of the model, we enforce non-negativity by applying a projection operator on *r*, [*Dp*]*e*, and *MD*1 at every Heun integration step.

#### 1.3.3 Network coupling of sub-networks

Regional inbound (i.e. afferent) drive for the three sub-networks was computed by applying their respective connectivity mask to the instantaneous firing-rate array **r**, with *w*^*e*^, *w*^*i*^, *w*^*d*^ serving as network-bound global coupling scalars:

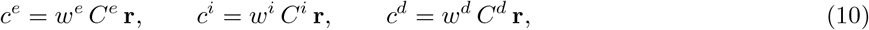

The left/right block structure of the connectome (resolved from per-node labels, due to the disparities in left/right block sizes across parcellations) serves solely to identify interhemispheric edges for subsequent feature computation in the cohort-amortized inference.

#### 1.3.4 Simulation and hemodynamic forward model

We integrated whole-brain dynamics through a stochastic Heun scheme with a time step of Δ*t* = 0.5 ms, and injected node-specific additive noise to the membrane-potential and adaptation equations using relative noise scalings of 0.1 and 0.01, respectively, with all other state variables remaining noise-free. Amortized simulations lasted a total of 450 s, with the initial 50 s discarded as transients. We executed simulations in a batched, multidevice parallel framework using JAX (Bradbury et al. 2018) (`jax.vmap` over the simulation batch within each device) so as to achieve the throughput required for the *N*sim = 20,000 simulations employed for training the density estimator (Section 1.5). We subsequently passed simulated regional firing-rate time series through a standard Balloon–Windkessel hemodynamic model (Buxton et al. 1998; Friston et al. 2000) to generate synthetic BOLD time series sampled at TR = 1 s.

#### 1.3.5 Fixed parameters of the whole-brain model

We present the biophysical constants governing local dynamics in Table 1, which underpin the equations in Eqs. 8–9 along with their associated firing-rate, membrane-potential, and adaptation formulations. We maintained these constants as identical across all nodes, with the sole exceptions of *Rd* and *I* (which we rendered heterogeneous across nodes as detailed in Section 1.3.1). We distinguish between two primary parameter groups. The top block of Table 1 describes population-scale and synaptic parameters following the previously established parameterization of Preti et al. (2026); Depannemaecker et al. (2026); Gast et al. (2024) and as utilized in Eqs. 8–9. The bottom block delineates network-scale parameters such as those revolving around external input, sub-network coupling, and dopaminergic-gain, which are primarily spatially agnostic global scalars, with *Rd, I* representing the only spatially informed exceptions noted above (Section 1.3.1). We re-calibrated this group’s values due to their original tuning stemming from the use of default-scale connectomes present in prior applications (including (Preti et al. 2026)). This deliberate choice was made as the log(1 + *w*)-harmonized connectomes employed throughout this study (Section 1.1.4) exhibit a systematically compressed dynamic range compared to raw structural connectomes.

**Table 1:** Parameters of the dopaminergic neural mass model. Top block: mean-field state-variable and synaptic constants, identical across nodes and unchanged from Preti et al. (2026). Bottom block: coupling, drive, and dopaminergic-gain parameters, re-tuned for the log-scale harmonized connectomes used in this study (see main text above); *I* and *Rd* are heterogeneous across nodes and reported as (baseline, elevated/scaled value).

| Parameter | Description | Value |
| --- | --- | --- |
| $C$ | Membrane capacitance scaling | 100.0 |
| $k$ | aQIF gain parameter | 0.7 |
| $\Delta$ | Half-width of the Lorentzian heterogeneity distribution | 0.5 |
| $v_r, v_\theta$ | Resting / threshold potential | −60.0, −40.0 |
| $B_d$ | Basal AMPA-conductance offset | 1.0 |
| $g_a, g_g$ | Baseline AMPA / GABA synaptic conductances | 1.0, 1.0 |
| $E_a, E_g$ | AMPA / GABA reversal potentials | 0.0, −80.0 |
| $\tau_{s_a}, \tau_{s_g}$ | AMPA / GABA synaptic time constants | 6.0, 6.0 |
| $\tau_u$ | Adaptation time constant | 33.33 |
| $b$ | Adaptation coupling to membrane potential | −2.0 |
| $\kappa$ | Adaptation coupling to firing rate | 100.0 |
| $\tau_{D_p}, \tau_{M_{D1}}$ | Dopamine concentration / D1-occupancy time constants | 500.0, 1000.0 |
| $K_m$ | Michaelis constant, DAT reuptake | 150.0 |
| $S_d$ | D1-occupancy sigmoid slope | −10.0 |
| $Z$ | D1-occupancy sigmoid log-offset | 0.5 |
| $\beta$ | D1-receptor sensitivity scaling | 1 |
| $w^e, w^i, w^d$ | Global excitatory / inhibitory / dopaminergic coupling | 0.12, 0.3, 0.03 |
| $\sigma$ | Global noise-amplitude scaling | 2 |
| $I$ | Tonic excitatory drive (baseline; bilateral pallidum) | 45.5; $45.5 \times 1.3$ |
| $J_a, J_g$ | Recurrent AMPA / GABA synaptic weights | 20.0, 0.0 |
| $J_{s_a}, J_{s_g}$ | Afferent AMPA / GABA synaptic weights | 0.08, 2.0 |
| $J_{D_p}$ | Afferent dopaminergic synaptic weight | 100000.0 |
| $R_d$ | Regional D1-receptor density | 0 outside $C^d$; $[0, 0.3]$ within $C^d$ |

### 1.4 Multimodal feature calculation

Empirical and simulated BOLD time series were reduced to a common set of structural and functional summary statistics following the excision of the initial transient. FC and functional connectivity dynamics (FCD) matrices were computed over the first 100 (cortical) nodes. Empirical BOLD time-series can possess acquisition-oriented and physiological noise, even after established de-noising procedures (Birn 2012). Furthermore, disparate coordination measures may result in significantly different connectivity patterns from the same source data (Liu et al. 2025). We therefore employed a shared measure of FC and, to the extent permitted by each source, a common FCD formulation across both empirical and synthetic sources, and subsequently harmonized the resulting summary statistics on a common scale as depicted in the feature-level harmonization of Section 1.1.4.

We first amplitude-normalized per-subject time-series by centering every region and dividing the entire array by the standard deviation pooled over regions and time. Normalizing time-series in this manner removes the arbitrary per-subject global signal amplitude whilst preserving inter-region amplitude variability. We subsequently computed functional connectivity via edge-level covariance (Liu et al. 2025; Strain et al. 2022) on these normalized time-series, identically across both empirical and synthetic data sources. Our FCD computation comprised vectorizing a time-resolved and source-specific connectivity pattern and Pearson-correlating the resulting vectors across time (Hansen et al. 2015). Due to the acquisition (i.e. different time-resolution) and duration disparities of the BOLD time series across cohorts, opting for a window of fixed length across all cohorts would impose a temporal scale varying with each cohort’s repetition time and run duration (Pedersen et al. 2018). Empirical FCD matrices were therefore computed via a frame-wise (rather than sliding-window) formulation, which equates to an instantaneous FC matrix calculated at every time point/frame as the cosine of the pairwise Hilbert phase difference, its upper triangle vectorized, with the resulting frame-by-frame arrays Pearson-correlated to yield the FCD matrix (Glerean et al. 2012).

Synthetic time series are, on the other hand, simulated under a fixed noise model and aim to represent the statistical temporal structure of BOLD dynamics (Kashyap and Keilholz 2019). Synthetic FCD was calculated via a sliding-window covariance procedure (window length *w* = 3 TR), wherein a covariance matrix is computed and vectorized per window and the window-by-window vectors are themselves Pearson-correlated to yield the FCD matrix, the variance of which is taken over the upper triangle at an offset of *w* windows, such that no pair of windows sharing a frame contributes to it. Windowing is indispensable on this source, as the per-frame noise of the simulated series would otherwise overwhelm an instantaneous estimate of coordination. Both approaches are established time-varying formulations of connectivity (Glerean et al. 2012; Pedersen et al. 2018). Similar to other summary statistics, we harmonized the interhemispheric FCD variances across sources prior to probabilistic inference (Section 1.1.4). We recomputed FCD matrices over a range of window sizes, which demonstrated an insensitivity of the chosen configuration to the window length. Recomputing over *w ∈ {*2, 3, 5, 7, 9*}* TR neither changed per-simulation ordering (Spearman *ρ ≥* 0.96 against *w* = 3 for 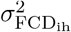, and *ρ ≥* 0.94 for 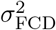), nor the monotone dependence upon *τ*_clear_ (*ρ* between *−*0.87 and *−*0.94 throughout). Similar to the functional connectomes, we restricted structural summary statistics to the first 100 (cortical) units and computed them directly from the corresponding block of the (harmonized) SC arrays, and ergo, the Harvard-Oxford subcortical nodes and the template-derived dopaminergic projection-source nodes (Section 1.3.1) do not contribute to the summary statistics employed in training the neural density estimator.

We selected a four-dimensional array (Section 1.5) as the minimal feature subset capable of jointly identifying ***θ***SC and *τ*_clear_ without redundancy (Section 1.4.1). Our functional features comprise 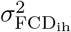 (variance of the upper-triangular FCD entries restricted to interhemispheric node pairs) and *σ*BOLD (BOLD-signal standard deviation; i.e. the temporal standard deviation of each region’s centered BOLD time series, averaged across the first 100 nodes). We computed *σ*BOLD prior to the per-subject rescaling applied to the connectivity estimates, to preserve the amplitude information which this feature aims to represent. The two structural features are 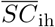 (mean interhemispheric connectome strength, cross-hemisphere and corpus-callosum-mediated) and ΔSC,intra (the signed intrahemispheric SC difference; i.e. the mean of the left-hemisphere intrahemispheric submatrix minus that of the right, the two submatrices node-matched to each other via homotopic correspondence). We also retained 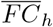(mean homotopic FC between each region and its contralateral homotope), along with global representations of FC/FCD 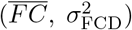, for exploratory comparisons. Finally, min–max normalization was applied per feature to all data features prior to training. Formally, we rescaled each summary statistic by its own minimum and maximum.

#### 1.4.1 Non-amortized parameter sweeps over *τ*_clear_

Prior to the configuration of the amortized inference, we validated the identifiability of the joint parameter space from a set of synthetic observations by executing two complementary sweeps. First, on a single reference connectome (that of the subject bearing the largest mean connectome strength), we simulated a 30 by 30 linearly spaced two-dimensional grid across *τ*_clear_ ([0.0001, 0.0025]) and an attenuation of only the interhemispheric edges of that connectome (intrahemispheric and dopaminergic-projection edges left intact), the latter traversing the full empirically observed span of mean interhemispheric strength (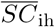 *∈* [0.11, 0.38] in harmonized, logtransformed units). Interhemispheric fluidity computed from these simulations reveals a non-degenerate mapping between *τ*_clear_ and 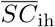 throughout the prior range, i.e. along the falling branch of the inverted-U (Fig. 2*e*). We additionally conducted a one-dimensional sweep over *τ*_clear_ (1,024 values over [0.0011, 0.0021]) for 7 subjects from the 1000BRAINS cohort, which were sampled evenly based on their 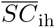 (Section 1.1.4). Complementing the previous 2D sweep, interhemispheric fluidity calculated from these simulations reveals non-degenerate and personalized feature-parameter curves which would be subsequently and faithfully represented in the amortized simulations (Fig. 2*f*).

### 1.5 Cohort-amortized inference of *τ*_clear_

ADAM’s joint amortized prior was constructed by computing the product of the CrossCoder-derived multi-variate normal over ***θ***SC (Section 1.2) and an independent uniform marginal prior over *τ*_clear_ *∈* [0.0011, 0.0020], yielding us the three-dimensional joint parameter space *{θ*_SC1_, *θ*_SC2_, *τ*_clear_*}*. We sampled from the joint prior (*N*sim = 20,000), and decoded the structural component of every sample through the decoder to generate a synthetic 118 *×* 118 connectome. These connectomes were subsequently fed as input to the whole-brain simulator (Section 1.3.4) with a shared set of heterogeneous baseline parameters, and partitioned by the fixed connectivity masks (Section 1.3.1). Simulated BOLD time series and previously decoded synthetic connectomes were then employed to attain the four-dimensional in-silico feature array (Section 1.4), thereby producing *N*sim distinct parameter-feature pairs.

We then fed synthetic parameter-feature arrays to an efficient class of neural density estimation known as neural spline flows (NSF), trained here in a single round as amortized neural posterior estimation (NPE; implemented by the `SNPE`class of the `sbi` package) (Durkan et al. 2019; Hashemi et al. 2023, 2024; Tejero-Cantero et al. 2020; Greenberg et al. 2019) to learn an invertible and efficient feature-parameter mapping. We eschewed further simulation rounds as amortization alleviates the need for (and matches the benefits reaped from) multi-round schemes. Empirical summary statistic (harmonized and normalized) arrays of the *N* = 3,650 participants were subsequently fed to the trained density estimator and we drew 1,000 posterior samples for each subject over *{θ*SC_1_, *θ*_SC2_, *τ*_clear_*}*. While full posterior distributions were retained to facilitate the age- and performance-stratified comparisons presented in Fig. 4*c, f, g*, we additionally computed per-dimension mean of each personalized array of posterior samples to arrive at a representative per-subject point estimate (e.g., Fig. 4*b, d, e*).

#### Structural moderation of Cog–τclear (Fig. 4g–i)

The moderating influence of structural integrity on the relationship between clearance timescales and cognitive performance was interrogated via a continuous interaction model (Section 1.1.5). Due to the much smaller scale of clearance timescale values, we chose to min–max normalize *τ*_clear_ to [0, 1] for this specific part of our investigation. Continuous analyses described below bear a mutual age-decorrelation approach (Fig. 4*g, h*). 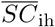 and Cog were regressed on a demeaned quadratic age polynomial, with their residuals SCresid and Cog_resid_ being z-scored and employed across both analyses.

We first fit the continuous interaction model once, 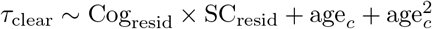 over the 36–95 age range. This interaction model yielded a single coefficient on Cog_resid_ *×* SCresid which constitutes our primary quantification of structure-dependence. We then employed a model-free shape assessment (Fig. 4*g*) that pools subjects across the full 36–95 year range, by partitioning SCresid into 6 bins and controlling for age_*c*_, performing intra-bin ordinary least-squares fit of *τ*_clear_ on Cog_resid_ aimed at recovering a local slope estimate. Subsequently, we leveraged a weighted and penalized natural-cubic spline (which employs knots at the bin centers and selects the smoothing parameter via leave-one-out cross-validation) to smooth said per-bin slopes and their associated standard errors, and conducted a parametric bootstrap over the estimated slopes so as to arrive at the pointwise 95% confidence band. The above process enables us to differentiate a genuine (and potentially asymmetric) crossover in the moderation from a mere artifact of extrapolating a stringently linear interaction term over the whole spectrum of connectome integrity.

Age-resolved conditional slopes were computed through a sliding-window approach. We partitioned subjects within each window (20-year width, 5-year step) into structurally preserved (SC-Pre) and structurally degraded (SC-Deg) subsets by the within-window median of age-residualized 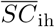, and fit *τ*_clear_ ~ Cog_resid_ + age_*c*_ independently for each partition via Huber-weighted robust regression (Fig. 4*h*). Windows with fewer than 300 subjects per partition were excluded from the analysis. Our step size of 5 years equates to substantial overlap of subjects between neighboring windows. In that light, we assessed significance with a cluster-based permutation approach (10^4^ age-stratified permutations, cluster-forming threshold |*t*| *>* 1.96, summed-|*t*| cluster mass), separately applied to the between-partition difference and to the SC-Deg coupling itself. We additionally fit 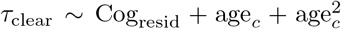 once over the full swept age range and for each partition’s subject set. This approach adopts a quadratic age term (as opposed to the linear term in the windowed approach), and aims to absorb *τ*_clear_’s curvature over the wider span. We refer to this non-overlapping fit as the per-partition pooled estimate, which serves as a multiplicity-free counterpart to the abovementioned windowed analysis.

We concluded this avenue of inquiry by conducting a twofold stratification of the inferred clearance timescale which can be viewed as a discrete portrait of the continuous interaction model (Fig. 4*i*). Guided by the continuous model’s findings, we constricted the comparison of personalized posteriors in high- versus low-cognitive performers to the 56–76 year age band (the range of centres of the significant run of SC-Deg sliding windows identified above). We define the strata quantifying structural degradation (namely SC-Pre vs SC-Deg), and those quantifying cognitive performance (namely, high-performers vs low-performers) via median splits of *SC*ih and Cog respectively (within the band, distinct from the windowed splits of age-residualized *SC*ih used above). Within this formulation, Cog is pooled across both SC strata so that the performance-tier threshold remains age-relative and common to both SC groups rather than being redefined separately within each.

Finally, we compared the groups using two-sided Brunner–Munzel tests, which are robust to unequal group sizes and unequal within-group variances, and calculated a rank-biserial effect size. In order to validate the robustness of significance values against the asymptotic test’s reference distribution, we permuted the performance labels within each degradation stratum (*n* = 2 *×* 10^4^), yielding a permutation significance value (*p*perm) for each stratum alongside three multiple-comparisons corrections. The tests comprise a Bonferroni correction, a Benjamini–Hochberg false-discovery-rate (*p*-value) permutation correction, and a Westfall–Young single-step max-*T* correction, respectively. The latter pools the largest test statistic across strata at every permutation draw into a joint null, and controls the family-wise error rate without assuming independence between strata (see Fig. 4*i* for the annotation of the max-*T p*-value).

### 1.6 Statistics and reproducibility

No statistical methods were used to predetermine sample sizes; sample sizes were instead set by the number of subjects per cohort with complete, quality-controlled diffusion, resting-state functional, and demographic/cognitive data surviving the exclusion criteria described in Sections 1.1 and 1.1.4. No additional data were excluded from any downstream analysis beyond those criteria. Because this study performs secondary, crosssectional analysis of previously collected observational neuroimaging cohorts rather than a controlled experiment, randomization and blinding of group allocation were not applicable; group membership (e.g., structurally degraded/preserved, high/low cognitive performer) was assigned post hoc from each subject’s own harmonized structural and cognitive measures. Where conducive, we report Pearson and Spearman correlations jointly (e.g., Fig. 1*h*–*m*). Two-sided *p*-values are computed from the standard Student’s-*t* approximation to the sampling distribution of the correlation coefficient under the null hypothesis of no association, 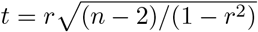 with *n −* 2 degrees of freedom. 10-year age-decade bins from 36–46 through 86–95 are used for the age-decade stratifications in Fig. 4*c* (sparse tail bins pooled where indicated). Equal-width bins spanning the range of Cog are employed for the cognitive-composite stratifications in Fig. 4*f*. All whole-brain simulations, CrossCoder training, and amortized density-estimator training and sampling were performed on a single workstation (Intel Core i9-14900K, 24 cores/32 threads; 128 GB RAM; one NVIDIA RTX 2000 Ada Generation GPU, 16 GB VRAM; Linux), without reliance on a shared computing cluster. All analyses were implemented in Python 3.12 using JAX 0.10 (whole-brain simulation and CrossCoder training), th`sbi` package (v0.26) for amortized neural posterior estimation, and `scipy` (v1.18) (Virtanen et al. 2020)/`statsmodels` (v0.14) (Seabold et al. 2010) for statistical testing; the complete software environment, with pinned package versions, is provided in the associated code repository.

This project/research has received funding from the European Union’s Horizon Europe Programme under the Specific Grant Agreement No. 101147319 (EBRAINS 2.0 Project), No. 101137289 (Virtual Brain Twin Project). Data, methods used, and/or research reported in this publication were provided in part by the Aging Adult Brain Connectome (AABC) project (U19AG073585) and the Human Connectome Project in Aging (HCP-A, U01AG052564) funded by the National Institute on Aging of the National Institutes of Health. HCP-A was further supported by funds provided by the McDonnell Center for Neuroscience at Washington University in St. Louis.

Data collection and sharing for this project was provided in part by the Cambridge Centre for Ageing and Neuroscience (CamCAN). CamCAN funding was provided by the UK Biotechnology and Biological Sciences Research Council (grant number BB/H008217/1), together with support from the UK Medical Research Council and University of Cambridge, UK.

Data collection and sharing for this project was provided in part by the 1000BRAINS study, which recruits from the Heinz Nixdorf Recall (HNR) Study and the HNR MultiGeneration Study. We thank the Heinz Nixdorf Foundation (Germany) for their generous support of the Heinz Nixdorf Recall Study. The HNR study is further supported by the German Ministry of Education and Science (FKZ 01EG9401) and the German Research Council (DFG, ER 155/6-1).

Data were provided in part by the enhanced Nathan Kline Institute-Rockland Sample (NKI-RS). Principal support for the enhanced NKI-RS project is provided by the NIMH BRAINS R01MH094639-01 (PI Milham). Funding for key personnel was also provided in part by the New York State Office of Mental Health and Research Foundation for Mental Hygiene.

Data were provided in part by OASIS-3: Principal Investigators: T. Benzinger, D. Marcus, J. Morris; NIH P30 AG066444, P50 AG00561, P30 NS09857781, P01 AG026276, P01 AG003991, R01 AG043434, UL1 TR000448, R01 EB009352.

## Author contributions

A.E.: Conceptualization, Data curation, Formal analysis, Investigation, Methodology, Software, Validation, Visualization, Writing – original draft, Writing – review & editing. G.P.: Methodology (initial dopaminergic model parameterization), Resources, Writing – review & editing. N.B., S.C.: Resources, Data curation (1000BRAINS), Writing – review & editing. M.M.W.: Methodology, Software (computational infrastructure), Writing – review & editing. M.H., S.P., D.M., V.J.: Writing – review & editing. All authors reviewed and approved the final manuscript.

## Data availability

The empirical neuroimaging and demographic data analyzed in this study originate from five previously published aging cohorts, all of which are available from their respective repositories/data access committees upon approval of an access request, subject to each consortium’s data-use terms: 1000BRAINS (Caspers et al. 2014), HCP-Aging (HCP-A) (Bookheimer et al. 2019), NKI-Rockland Sample (NKI-RS) (Nooner et al. 2012), Cam-CAN (Shafto et al. 2014; Taylor et al. 2017), and OASIS-3 (LaMontagne et al. 2019). Empirically observed PET/SPECT dopamine-transporter data underlying Fig. 1*b* (Methods, Section 1.1.6) are openly available from Karrer et al. (2017) at osf.io/6uf82. The NKI-RS neuroimaging releases are accessible via the project’s current data-release portal at rocklandsample.org/accessing-the-neuroimaging-data-releases. A Schaefer 100-parcel structural/functional connectivity release for the 1000BRAINS cohort is further available via the EBRAINS Knowledge Graph at search.kg.ebrains.eu/instances/3f179784-194d-4795-9d8d-301b524ca00a. The trained CrossCoder, alongside synthetic connectomes generated thereof, and amortized density-estimator model artifacts, will be made publicly available upon publication of this work. Subject-level empirical connectomes and personalized posterior samples are not released, so as to preclude re-identification of individual participants from connectome-derived representations (cf. the subject-identifiability results of Fig. 2*j*).

## Code availability

All simulation, whole-brain model, CrossCoder, amortized inference, posterior sampling, and statistical-analysis code will be made publicly available, with a versioned release and DOI (e.g. via Zenodo), upon publication of this work, at gitlab.ebrains.eu/ahesmaeili/adam.

## References

Abraham A, Pedregosa F, Eickenberg M, et al (2014) Machine learning for neuroimaging with scikit-learn. Frontiers in Neuroinformatics 8:14. doi: 10.3389/fninf.2014.00014

Battaglia D, Boudou T, Hansen ECA, et al (2020) Dynamic functional connectivity between order and randomness and its evolution across the human adult lifespan. NeuroImage 222:117156. doi: 10.1016/j.neuroimage.2020.117156

Behrens TEJ, Berg HJ, Jbabdi S, et al (2007) Probabilistic diffusion tractography with multiple fibre orientations: What can we gain? NeuroImage 34(1):144–155. doi: 10.1016/j.neuroimage.2006.09.018

Bethlehem RA, Seidlitz J, White SR, et al (2022) Brain charts for the human lifespan. Nature 604(7906):525–533. doi: 10.1038/s41586-022-04554-y

Birn RM (2012) The role of physiological noise in resting-state functional connectivity. NeuroImage 62(2):864– 870. doi: 10.1016/j.neuroimage.2012.01.016

Björklund A, Dunnett SB (2007) Dopamine neuron systems in the brain: an update. Trends in Neurosciences 30(5):194–202. doi: 10.1016/j.tins.2007.03.006

Bookheimer SY, Salat DH, Terpstra M, et al (2019) The lifespan human connectome project in aging: an overview. NeuroImage 185:335–348. doi: 10.1016/j.neuroimage.2018.10.009

Bradbury J, Frostig R, Hawkins P, et al (2018) JAX: composable transformations of Python+NumPy programs. URL http://github.com/jax-ml/jax

Buxton RB, Wong EC, Frank LR (1998) Dynamics of blood flow and oxygenation changes during brain activation: the balloon model. Magnetic Resonance in Medicine 39(6):855–864. doi: 10.1002/mrm.1910390602

Caspers S, Moebus S, Lux S, et al (2014) Studying variability in human brain aging in a population-based German cohort—rationale and design of 1000BRAINS. Frontiers in Aging Neuroscience 6:149. doi: 10.3389/fnagi.2014.00149

Chan MY, Park DC, Savalia NK, et al (2014) Decreased segregation of brain systems across the healthy adult lifespan. Proceedings of the National Academy of Sciences 111(46):E4997–E5006. doi: 10.1073/pnas.1415122111

Cong J, Zhao S, Xu H, et al (2026) White-matter functional connectivity uniquely predicts brain age, cognition, and psychopathology beyond gray-matter connectivity. Proceedings of the National Academy of Sciences 123(34):e2604191123. doi: 10.1073/pnas.2604191123

Cools R, D’Esposito M (2011) Inverted-u-shaped dopamine actions on human working memory and cognitive control. Biological Psychiatry 69(12):e113–e125. doi: 10.1016/j.biopsych.2011.03.028

Depannemaecker D, Duprat C, Casagrande G, et al (2026) A next generation neural mass model with neuromodulation. Journal of Computational Neuroscience 54(1):23–43. doi: 10.1007/s10827-026-00920-1

Dreher JC, Burnod Y (2002) An integrative theory of the phasic and tonic modes of dopamine modulation in the prefrontal cortex. Neural Networks 15(4–6):583–602. doi: 10.1016/S0893-6080(02)00051-5

Durkan C, Bekasov A, Murray I, et al (2019) Neural spline flows. In: Advances in Neural Information Processing Systems, vol 32. Curran Associates, Inc., URL https://proceedings.neurips.cc/paper_files/paper/2019/file/7ac71d433f282034e088473244df8c02-Paper.pdf

Esmaeili A, Woodman MM, Baldy N, et al (2026) Cohort-amortized personalization: navigating the privacy-utility frontier for virtual brain twins. arXiv doi: 10.48550/arXiv.2606.30329

Esteban O, Markiewicz CJ, Blair RW, et al (2019) fMRIPrep: a robust preprocessing pipeline for functional MRI. Nature Methods 16(1):111–116. doi: 10.1038/s41592-018-0235-4

Frazier JA, Chiu S, Breeze JL, et al (2005) Structural brain magnetic resonance imaging of limbic and thalamic volumes in pediatric bipolar disorder. American Journal of Psychiatry 162(7):1256–1265. doi: 10.1176/appi.ajp.162.7.1256

Friston KJ, Williams S, Howard R, et al (1996) Movement-related effects in fMRI time-series. Magnetic Resonance in Medicine 35(3):346–355. doi: 10.1002/mrm.1910350312

Friston KJ, Mechelli A, Turner R, et al (2000) Nonlinear responses in fmri: the balloon model, volterra kernels, and other hemodynamics. NeuroImage 12(4):466–477. doi: 10.1006/nimg.2000.0630

Gast R, Solla SA, Kennedy A (2024) Neural heterogeneity controls computations in spiking neural networks. Proceedings of the National Academy of Sciences 121(3):e2311885121. doi: 10.1073/pnas.2311885121

Glerean E, Salmi J, Lahnakoski JM, et al (2012) Functional magnetic resonance imaging phase synchronization as a measure of dynamic functional connectivity. Brain Connectivity 2(2):91–101. doi: 10.1089/brain.2011.0068

Goldstein JM, Seidman LJ, Makris N, et al (2007) Hypothalamic abnormalities in schizophrenia: sex effects and genetic vulnerability. Biological Psychiatry 61(8):935–945. doi: 10.1016/j.biopsych.2006.06.027

Greenberg D, Nonnenmacher M, Macke J (2019) Automatic posterior transformation for likelihood-free inference. In: International conference on machine learning, PMLR, pp 2404–2414

Hansen ECA, Battaglia D, Spiegler A, et al (2015) Functional connectivity dynamics: Modeling the switching behavior of the resting state. NeuroImage 105:525–535. doi: 10.1016/j.neuroimage.2014.11.001

Harms MP, Somerville LH, Ances BM, et al (2018) Extending the human connectome project across ages: imaging protocols for the lifespan development and aging projects. NeuroImage 183:972–984. doi: 10.1016/j.neuroimage.2018.09.060

Hashemi M, Vattikonda AN, Jha J, et al (2023) Amortized Bayesian inference on generative dynamical network models of epilepsy using deep neural density estimators. Neural Networks 163:178–194. doi: 10.1016/j.neunet.2023.03.040

Hashemi M, Ziaeemehr A, Woodman MM, et al (2024) Simulation-based inference on virtual brain models of disorders. Machine Learning: Science and Technology 5(3):035019. doi: 10.1088/2632-2153/ad6230

Hashemi M, Depannemaecker D, Saggio M, et al (2025) Principles and operation of virtual brain twins. IEEE Reviews in Biomedical Engineering 19:111–139. doi:10.1109/RBME.2025.3562951

Hunger L, Kumar A, Schmidt R (2020) Abundance compensates kinetics: similar effect of dopamine signals on D1 and D2 receptor populations. Journal of Neuroscience 40(14):2868–2881. doi: 10.1523/JNEUROSCI.1951-19.2019

Jenkinson M, Beckmann CF, Behrens TEJ, et al (2012) Fsl. NeuroImage 62(2):782–790. doi: 10.1016/j.neuroimage.2011.09.015

Kaller S, Rullmann M, Patt M, et al (2017) Test–retest measurements of dopamine d1-type receptors using simultaneous pet/mri imaging. European journal of nuclear medicine and molecular imaging 44(6):1025–1032. doi: 10.1007/s00259-017-3645-0

Kanel P, Koeppe RA, Kotagal V, et al (2023) Regional serotonin terminal density in aging human brain: a [11c] dasb pet study. Aging Brain 3:100071. doi: 10.1016/j.nbas.2023.100071

Karrer TM, Josef AK, Mata R, et al (2017) Reduced dopamine receptors and transporters but not synthesis capacity in normal aging adults: a meta-analysis. Neurobiology of Aging 57:36–46. doi: 10.1016/j.neurobiolaging.2017.05.006

Kashyap A, Keilholz S (2019) Dynamic properties of simulated brain network models and empirical resting-state data. Network Neuroscience 3(2):405–426. doi: 10.1162/netn_a_00070

Korkki SM, Johansson J, Nordin K, et al (2025) Dedifferentiation of caudate functional organization is linked to reduced d1 dopamine receptor availability and poorer memory function in aging. Imaging Neuroscience 3:1–20. doi: 10.1162/imag_a_00462

Kroener S, Chandler LJ, Phillips PEM, et al (2009) Dopamine modulates persistent synaptic activity and enhances the signal-to-noise ratio in the prefrontal cortex. PLoS ONE 4(8):e6507. doi: 10.1371/journal.pone.0006507

Lalwani P, Gagnon H, Cassady K, et al (2019) Neural distinctiveness declines with age in auditory cortex and is associated with auditory gaba levels. NeuroImage 201:116033. doi: 10.1016/j.neuroimage.2019.116033

LaMontagne PJ, Benzinger TLS, Morris JC, et al (2019) OASIS-3: Longitudinal neuroimaging, clinical, and cognitive dataset for normal aging and Alzheimer disease. medRxiv doi: 10.1101/2019.12.13.19014902

Lanciego JL, Luquin N, Obeso JA (2012) Functional neuroanatomy of the basal ganglia. Cold Spring Harbor perspectives in medicine 2(12):a009621. doi: 10.1101/cshperspect.a009621

Lavanga M, Stumme J, Yalcinkaya BH, et al (2023) The virtual aging brain: Causal inference supports interhemispheric dedifferentiation in healthy aging. NeuroImage 283:120403. doi: 10.1016/j.neuroimage.2023.120403

Li SC, Rieckmann A (2014) Neuromodulation and aging: implications of aging neuronal gain control on cognition. Current Opinion in Neurobiology 29:148–158. doi: 10.1016/j.conb.2014.07.009

Li SC, Lindenberger U, Sikström S (2001) Aging cognition: from neuromodulation to representation. Trends in cognitive sciences 5(11):479–486. doi: 10.1016/S1364-6613(00)01769-1

Liu ZQ, Luppi AI, Hansen JY, et al (2025) Benchmarking methods for mapping functional connectivity in the brain. Nature Methods 22(7):1593–1602. doi: 10.1038/s41592-025-02704-4

Makris N, Goldstein JM, Kennedy D, et al (2006) Decreased volume of left and total anterior insular lobule in schizophrenia. Schizophrenia Research 83(2-3):155–171. doi: 10.1016/j.schres.2005.11.020

Marcus DS, Harwell J, Olsen T, et al (2011) Informatics and data mining tools and strategies for the human connectome project. Frontiers in Neuroinformatics 5:4. doi: 10.3389/fninf.2011.00004

Mollink J, Smith SM, Elliott LT, et al (2019) The spatial correspondence and genetic influence of interhemispheric connectivity with white matter microstructure. Nature neuroscience 22(5):809–819. doi: 10.1038/s41593-019-0379-2

Mueller A, Krock RM, Shepard S, et al (2020) Dopamine receptor expression among local and visual cortex-projecting frontal eye field neurons. Cerebral Cortex 30(1):148–164. doi: 10.1093/cercor/bhz078

Nicholson C (1995) Interaction between diffusion and michaelis-menten uptake of dopamine after iontophoresis in striatum. Biophysical Journal 68(5):1699–1715. doi: 10.1016/S0006-3495(95)80348-6

Nooner KB, Colcombe SJ, Tobe RH, et al (2012) The NKI-Rockland sample: a model for accelerating the pace of discovery science in psychiatry. Frontiers in Neuroscience 6:152. doi: 10.3389/fnins.2012.00152

Pedersen M, Omidvarnia A, Zalesky A, et al (2018) On the relationship between instantaneous phase synchrony and correlation-based sliding windows for time-resolved fmri connectivity analysis. NeuroImage 181:85–94. doi: 10.1016/j.neuroimage.2018.06.020

Pedersen R, Geerligs L, Andersson M, et al (2021) When functional blurring becomes deleterious: Reduced system segregation is associated with less white matter integrity and cognitive decline in aging. NeuroImage 242:118449. doi: 10.1016/j.neuroimage.2021.118449

Pedersen R, Johansson J, Salami A (2023) Dopamine d1-signaling modulates maintenance of functional network segregation in aging. Aging Brain 3:100079. doi: 10.1016/j.nbas.2023.100079

Petkoski S, Ritter P, Jirsa VK (2023) White-matter degradation and dynamical compensation support age-related functional alterations in human brain. Cerebral Cortex 33(10):6241–6256. doi: 10.1093/cercor/bhac500

Power JD, Barnes KA, Snyder AZ, et al (2012) Spurious but systematic correlations in functional connectivity MRI networks arise from subject motion. NeuroImage 59(3):2142–2154. doi: 10.1016/j.neuroimage.2011.10.018

Power JD, Mitra A, Laumann TO, et al (2014) Methods to detect, characterize, and remove motion artifact in resting state fMRI. NeuroImage 84:320–341. doi: 10.1016/j.neuroimage.2013.08.048

Preti G, Casagrande G, Wang H, et al (2025) A neural mass model with dopaminergic d1 neuromodulation (0.1.0). doi: 10.25493/9KR4-BNK

Preti G, Wang HE, Ziaeemehr A, et al (2026) Virtual brain twins guide personalized treatment decision in schizophrenia. medRxiv doi: 10.64898/2026.05.06.26352533

Puxeddu MG, Faskowitz J, Betzel RF, et al (2020) The modular organization of brain cortical connectivity across the human lifespan. NeuroImage 218:116974. doi: 10.1016/j.neuroimage.2020.116974

Roland JL, Snyder AZ, Hacker CD, et al (2017) On the role of the corpus callosum in interhemispheric functional connectivity in humans. Proceedings of the National Academy of Sciences 114(50):13278–13283. doi: 10.1073/pnas.1707050114

Schaefer A, Kong R, Gordon EM, et al (2018) Local-global parcellation of the human cerebral cortex from intrinsic functional connectivity MRI. Cerebral Cortex 28(9):3095–3114. doi: 10.1093/cercor/bhx179

Seabold S, Perktold J, et al (2010) Statsmodels: econometric and statistical modeling with python. scipy 7(1):92– 96

Shafto MA, Tyler LK, Dixon M, et al (2014) The Cambridge Centre for Ageing and Neuroscience (Cam-CAN) study protocol: a cross-sectional, lifespan, multidisciplinary examination of healthy cognitive ageing. BMC Neurology 14:204. doi: 10.1186/s12883-014-0204-1

Strain JF, Brier MR, Tanenbaum A, et al (2022) Covariance-based vs. correlation-based functional connectivity dissociates healthy aging from alzheimer disease. NeuroImage 261:119511. doi: 10.1016/j.neuroimage.2022.119511

Stumme J, Krämer C, Miller T, et al (2022) Interrelating differences in structural and functional connectivity in the older adult’s brain. Human Brain Mapping 43(18):5543–5561. doi: 10.1002/hbm.26030

Sullivan EV, Rohlfing T, Pfefferbaum A (2010) Quantitative fiber tracking of lateral and interhemispheric white matter systems in normal aging: Relations to timed performance. Neurobiology of Aging 31(3):464–481. doi: 10.1016/j.neurobiolaging.2008.04.007

Sun X, Zhao Y, Wolf ME (2005) Dopamine receptor stimulation modulates AMPA receptor synaptic insertion in prefrontal cortex neurons. Journal of Neuroscience 25(32):7342–7351. doi:10.1523/JNEUROSCI.4603-04.2005

Taylor JR, Williams N, Cusack R, et al (2017) The cambridge centre for ageing and neuroscience (cam-can) data repository: Structural and functional mri, meg, and cognitive data from a cross-sectional adult lifespan sample. neuroimage 144:262–269. doi: 10.1016/j.neuroimage.2015.09.018

Tejero-Cantero A, Boelts J, Deistler M, et al (2020) sbi: A toolkit for simulation-based inference. Journal of Open Source Software 5(52):2505. doi: 10.21105/joss.02505

Thurley K, Senn W, Lüscher HR (2008) Dopamine increases the gain of the input–output response of rat prefrontal pyramidal neurons. Journal of Neurophysiology 99(6):2985–2997. doi: 10.1152/jn.01098.2007

Tournier JD, Smith R, Raffelt D, et al (2019) MRtrix3: A fast, flexible and open software framework for medical image processing and visualisation. NeuroImage 202:116137. doi: 10.1016/j.neuroimage.2019.116137

Virtanen P, Gommers R, Oliphant TE, et al (2020) SciPy 1.0: Fundamental Algorithms for Scientific Computing in Python. Nature Methods 17:261–272. doi: 10.1038/s41592-019-0686-2

Westlye LT, Walhovd KB, Dale AM, et al (2010) Life-span changes of the human brain white matter: Diffusion tensor imaging (dti) and volumetry. Cerebral Cortex 20(9):2055–2068. doi: 10.1093/cercor/bhp280

Zhang H, Cao P, Mak HKF, et al (2024) The structural–functional-connectivity coupling of the aging brain. GeroScience 46(4):3875–3887. doi: 10.1007/s11357-024-01106-2

Zhang Y, Larcher KMH, Misic B, et al (2017) Anatomical and functional organization of the human substantia nigra and its connections. elife 6:e26653. doi: 10.7554/eLife.26653

