## Supplementary Information for "Structural integrity moderates the cognitive bearing of dopamine clearance in aging"

Figure, table, and section numbers prefixed with **S** refer to this document. All other figure, table, section, and equation numbers refer to the main article.

#### Contents


| No. | Section | Page |
|---|---|---|
| S1. | Synthetic validation, posterior contraction, and null-hypothesis testing (Fig. S1) | 2 |
| S2. | CrossCoder architecture selection across latent dimensionalities (Fig. S2) | 6 |
| S3. | Synthetic parameter-feature mappings (Fig. S3) | 6 |
| S4. | Node-resolved simulations under a diffuse prior of clearance timescales (Fig. S6) | 8 |
| S5. | Empirical summary statistics versus age (Fig. S4) | 9 |
| S6. | Subject-level structural connectome reconstruction exemplars (Fig. S5) | 10 |
| S7. | Per-task validation of the cognitive composite across cohorts (Fig. S7) | 10 |
| S8. | Disentangling the influence of structural decay versus dopamine clearance timescales on cognitive performance (Table S2) | 13 |
| S9. | Cognitive tasks grouped by domain (Table S4) | 14 |

### 1 Supplementary Information

#### S1. Synthetic validation, posterior contraction, and null-hypothesis testing

Five complementary checks were employed to ensure the informativeness of ADAM's four-dimensional feature vector $\{\sigma^2_{\mathrm{FCD_{ih}}}, \sigma_{\mathrm{BOLD}}, \overline{SC}_{\mathrm{ih}}, \Delta_{\mathrm{SC,intra}}\}$ on the amortized posterior, compared to the more diffuse joint prior. Our audit substantiates the ensuing age-associations observed in $\theta_{\mathrm{SC}_1}, \theta_{\mathrm{SC}_2}$, and $\tau_{\mathrm{clear}}$ surpassing the magnitude anticipated from exclusively random feature-subject pairings. The fourth check directly assesses whether the CrossCoder's structural compression itself distorts the resulting posterior. The fifth check involves quantification of parameter recovery and the calibration of the posterior's credible intervals.

Held-out synthetic validation serves as one exemplar of the above informativeness (Supplementary Fig. S1**a**): where we observe not only high posterior shrinkage, but also a robust convergence of the 'predicted' posterior means (or peaks in multimodal sampled distributions) towards the 'observed' parameter values. This convergence was quantified via computation of an uncertainty-reduction measure (i.e. posterior shrinkage; $1 - \sigma_{\mathrm{post}}/\sigma_{\mathrm{prior}}$) directly from posterior samples over 150 training simulation trials (300 draws per observable; Supplementary Fig. S1**b**). Uncertainty reduction comprised $\approx$ 0.90 for $\theta_{\mathrm{SC}_1}$, $\approx$ 0.99 for $\theta_{\mathrm{SC}_2}$, and $\approx$ 0.84 for $\tau_{\mathrm{clear}}$ (each echoing the mapping onto their representative summary statistic, as alluded to in Fig. S3). This computation was subsequently replicated on empirical observations, which reproduced the same ordinal hierarchy (median $\approx 0.91$, $\approx 0.85$, and $\approx 0.99$ for $\theta_{\mathrm{SC}_1}, \tau_{\mathrm{clear}}$, and $\theta_{\mathrm{SC}_2}$, respectively).

We initially explored the native active-subspace projection of `sbi` as a possible proxy for our sample-oriented approach, but abandoned it due to its systemic inaptness to robustly capture the informativeness of $\tau_{\mathrm{clear}}$. The prior distribution of the clearance timescales occupies a narrow uniform interval, compared to the wide multivariate normal structural priors. Accordingly, the sensitivity this projection attributes to $\tau_{\mathrm{clear}}$ ($\approx 0.2\%$ of the total normalized sensitivity, against $\approx 48 - 52\%$ for $\theta_{SC}$) stands at odds with its substantial posterior contraction ($\approx 85\%$ uncertainty reduction). Sensitivity scores derived from Fisher information (tuned to the local curvature of log-posterior) thus largely fail to capture the informativeness of a parameter sampled from a comparatively constricted prior, which we addressed by derivation of a data-centric proxy. To that end, we employed the first two principal components of the gathered, $z$-standardized empirical posterior samples (accounting for $\approx$54% and $\approx$34% of the posterior variance, respectively), with our derivation stemming solely from the inherent sample dispersion as opposed to local gradients (Fig. S1**c**). Serving as an exemplar, a representative empirical subject with the nearest posterior mean to the centroid of the aggregate posterior means (in $z$-scored parameter space) was projected onto this basis, providing further support for the substantial posterior shrinkage under said joint projection (as opposed to the uninformative gradient-based metrics).

We additionally conducted a posterior predictive check to directly assess whether CrossCoder compression of connectomes influences the $\tau_{\mathrm{clear}}$ read-out. This was done by resimulating the macroscale model for $n = 500$ subjects once using each subject's own uncompressed SC at that subject's amortized posterior estimation of $\tau_{\mathrm{clear}}$ (i.e. posterior mean), computing a similar feature array and subsequent feeding said array back through the previously trained density estimator. We found that the $\tau_{\mathrm{clear}}$ readout from simulations done using uncompressed-connectome correlated strongly with the original amortized (decoded-SC) estimates (Pearson $r = 0.85$, Spearman $\rho = 0.85$, both $p < 10^{-138}$; Supplementary Fig. S1**d**), further supporting the notion that CrossCoder connectome compression does not meaningfully distort the posterior estimates of clearance timescales compared to direct employment of subjects' uncompressed connectomes.

Finally, we tested whether posterior means can recover simulated ground-truth and whether the posterior's credible intervals are correctly sized. Our analysis revealed that posterior means accurately track true values for both the structural and neuromodulatory parameters (across 5,000 of the $N_{\mathrm{sim}} = 20{,}000$ training simulations confined to the empirical feature envelope; Pearson $r \geq 0.9999$ for $\theta_{\mathrm{SC}_1}$ and $\theta_{\mathrm{SC}_2}$, and $r = 0.982$ for $\tau_{\mathrm{clear}}$; Supplementary Fig. S1**e–g**). We additionally assessed calibration through the TARP expected-coverage diagnostic, which was near-exact under both the global prior and the empirical envelope (area-to-calibration $-0.002$ and $+0.0002$; Kolmogorov–Smirnov $p > 0.99$ for both; Supplementary Fig. S1**h**). Taken together, these results suggest that the amortized posterior tracks simulated parameters closely and is not overconfident within the regime the empirical data occupy.

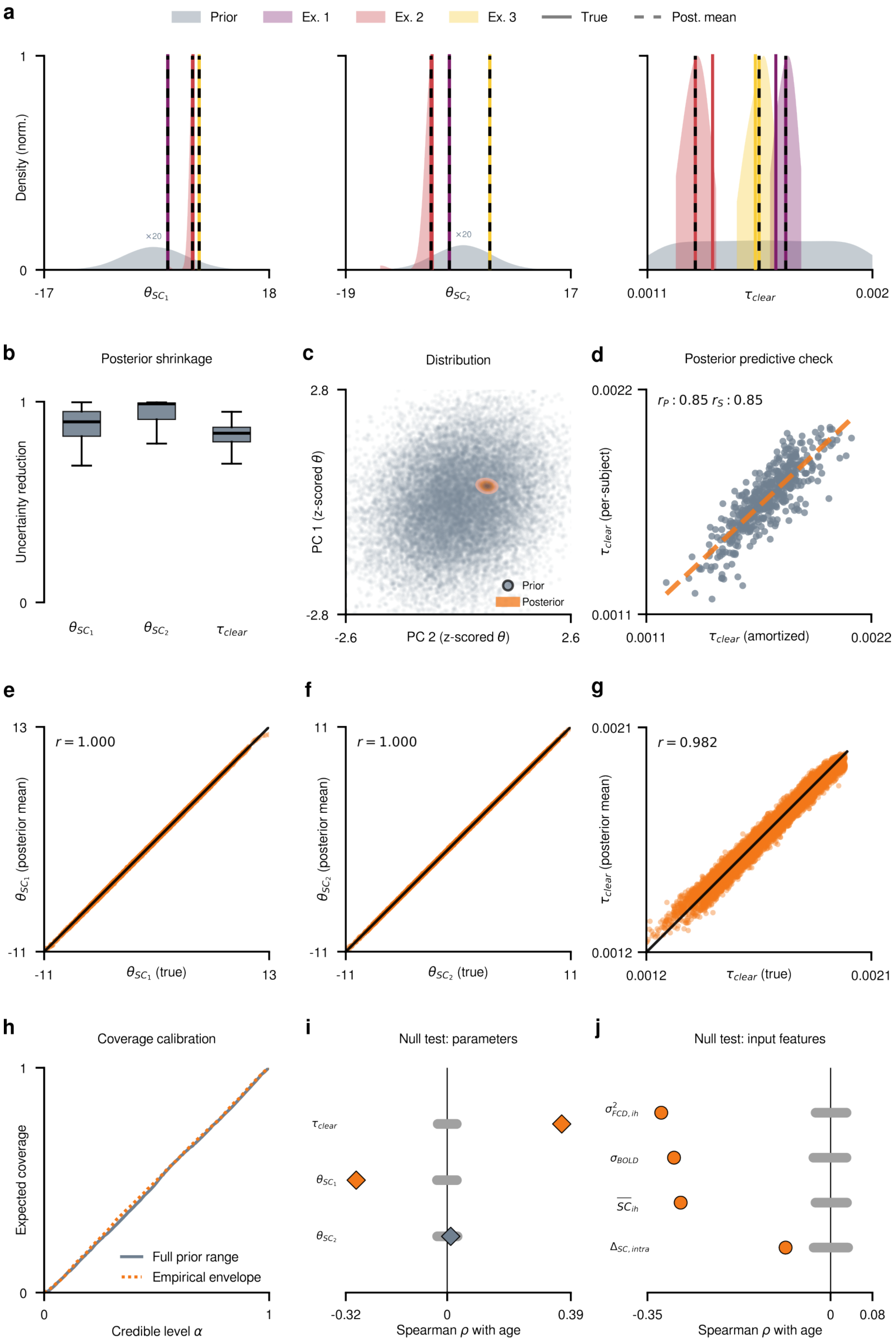


**Fig. S1**: **Synthetic validation, posterior contraction, and null-hypothesis testing of the amortized posterior. a,** Prior (grey) and posterior densities for three representative training simulations (Ex. 1–3), for each of the three inferred parameters $\theta_{\mathrm{SC}_1}$, $\theta_{\mathrm{SC}_2}$, and $\tau_{\mathrm{clear}}$; solid vertical lines mark the true simulated value, dashed lines the posterior mean. Posteriors are drawn at unit peak height and the prior at its true ratio to the sharpest of them, which for the structural parameters leaves it too flat to see; both are therefore drawn magnified $\times 20$ (annotated in-panel), whereas the prior of $\tau_{\mathrm{clear}}$ is drawn unmagnified. The magnification is a legibility choice alone, and conveys how sharply each posterior contracts relative to its prior. **b,** Posterior shrinkage $(1 - \sigma_{\mathrm{post}}/\sigma_{\mathrm{prior}})$ across 150 training simulations (300 posterior samples each), shown as boxplots to capture the occasionally bimodal shape of the $\tau_{\mathrm{clear}}$ posterior. **c,** Posterior distribution for a representative empirical subject, projected onto the top two principal components of the pooled, z-scored empirical posterior samples (grey: prior samples; orange: posterior, kernel density estimate). **d,** Posterior predictive check: estimated $\tau_{\mathrm{clear}}$ for $n = 500$ from cohort-amortization pipeline versus $\tau_{\mathrm{clear}}$ re-estimation by simulating each subject's own uncompressed SC at that subject's amortized posterior-mean $\tau_{\mathrm{clear}}$ (and subsequent feeding of the resulting features back through the same trained estimator; $r = 0.85$, $\rho_S = 0.85$, both $p < 10^{-138}$). **e–g,** Parameter recovery over 5,000 of the $N_{\mathrm{sim}} = 20{,}000$ training simulations drawn within the empirical feature envelope (300 posterior samples each): posterior mean against the true simulated value for $\theta_{\mathrm{SC}_1}$ (**e**), $\theta_{\mathrm{SC}_2}$ (**f**), and $\tau_{\mathrm{clear}}$ (**g**); Pearson $r \geq 0.9999$ for both structural parameters and $r = 0.982$ for $\tau_{\mathrm{clear}}$. **h,** TARP expected coverage against credibility level, with the global-prior and empirical-envelope scopes overlaid; the diagonal marks exact calibration (area-to-calibration $-0.002$ and $+0.0002$ respectively, Kolmogorov–Smirnov $p > 0.99$ for both). **i,** Null-hypothesis testing of all estimated parameters: observed Spearman correlation with age (colour-coded markers) against the null distribution obtained by permuting the feature-to-subject assignment and re-inferring posteriors from the trained density estimator (500 repetitions per parameter), ranked by $|\rho|$; grey segments show each parameter's 95% null interval. **j,** As in **i**, for the four input features entering the observation model, using a simple age-shuffle null (no posterior resampling required for raw features). Orange markers in **i–j** indicate observables falling outside their null interval.

Null-hypothesis testing was conducted by permutation of the observables and their subsequent re-inference ($n_{iter} = 500$). For each iteration we applied a shared random permutation jointly to all four dimensions of the empirical feature array, which preserves each subject's observables while shuffling their metadata assignment.

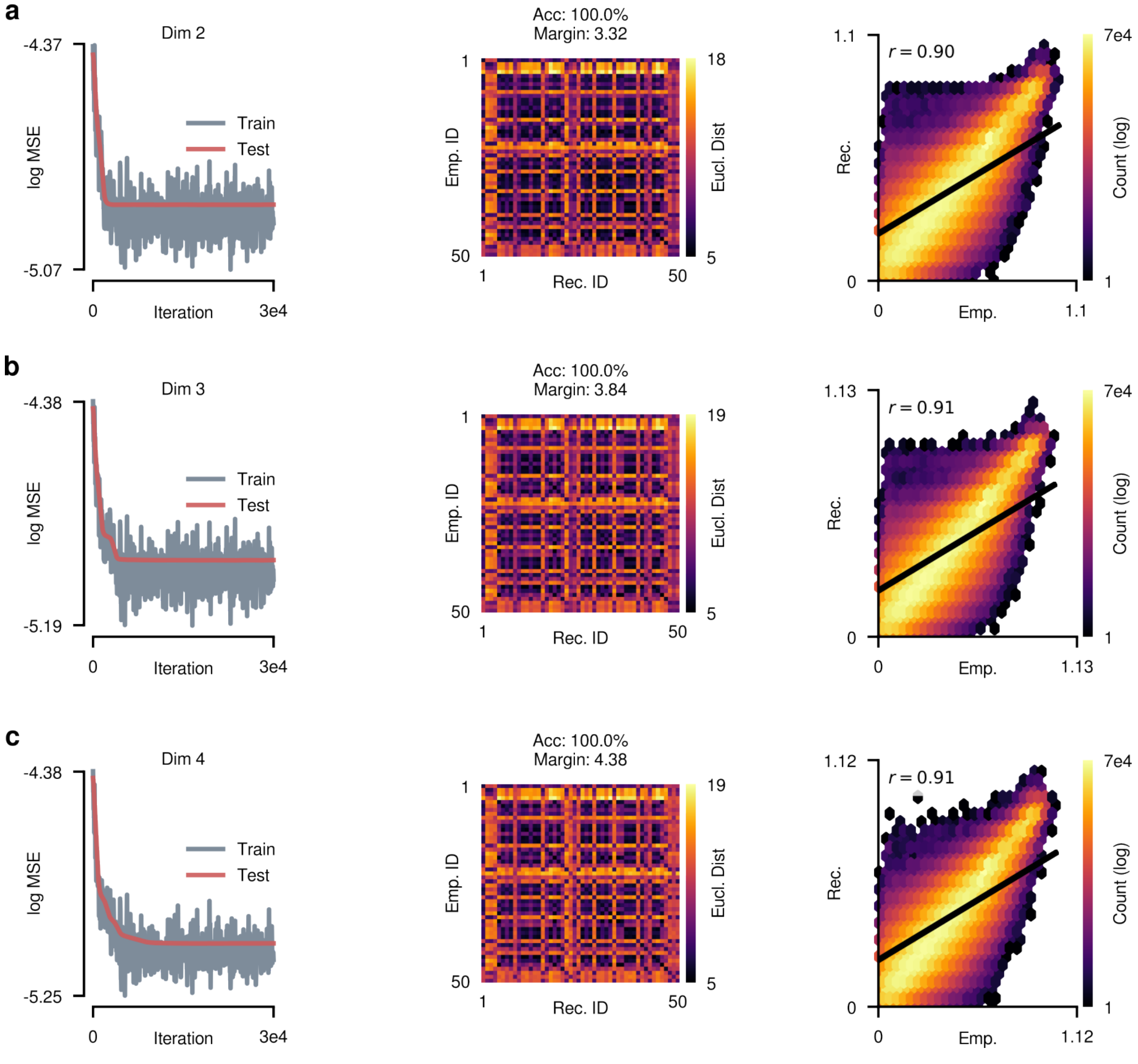


**Fig. S2**: **CrossCoder architecture selection across candidate latent dimensionalities. a–c,** Training (grey) and test (salmon) reconstruction-MSE traces (left), held-out subject-identifiability confusion matrices with nearest-reconstruction classification accuracy (middle), and empirical-vs-reconstructed SC edge-weight fidelity (right, Spearman $\rho$), for CrossCoder latent dimensionality $n_{\mathrm{lat}} = 2$ (**a**, the architecture retained throughout the main text), $n_{\mathrm{lat}} = 3$ (**b**), and $n_{\mathrm{lat}} = 4$ (**c**).

We then re-interrogated the trained density estimator (via batched sampling) on said permuted data to generate a robust set of inferred null samples with respect to each parameter (as opposed to a mere age-shuffling of previously estimated samples). Starting with the clearance timescales, we noted that the observed Spearman correlation with age ($\rho = +0.36$, computed from the fresh 300-sample posterior draw used for this test; $+0.37$ for the 1,000-sample posterior means of main-text Fig. 4*d*) dwelled far beyond the null distribution (95% interval

$\approx [-0.03, +0.03]$), with no single re-inferred iteration producing an association of comparable magnitude with an empirical $p < 1/500$ (Supplementary Fig. S1**i**). Our findings reveal that all observables and two out of three parameters dwell beyond their respective empirical 95% null distribution. This provides further evidence for the previously reported age-associations, as opposed to their stemming from a particular choice of prior range or stochastic metadata-feature pairing ($\rho = -0.29$, $\rho = +0.36$, $\rho \approx -0.32$, $\rho \approx -0.30$, $\rho \approx -0.29$, $\rho \approx -0.09$ for $\theta_{\mathrm{SC}_1}$, $\tau_{\mathrm{clear}}$, $\sigma^2_{\mathrm{FCD_{ih}}}$, $\sigma_{\mathrm{BOLD}}$, $\overline{SC}_{\mathrm{ih}}$, and $\Delta_{\mathrm{SC,intra}}$, respectively; Supplementary Fig. S1**i,j**). The four observables were tested by a direct age-shuffling of summary statistics, rather than by the re-inference procedure described above for the estimated parameters. The only exception lies with $\theta_{\mathrm{SC}_2}$ ($\rho = +0.01$), which one could anticipate from the weak age-association found within its corresponding observable. Put another way, $\Delta_{\mathrm{SC,intra}}$ itself exhibits a faint and inconsistent cross-cohort trend, with the pooling effect primarily inducing this marginal crossing of non-significance. Aiming to corroborate this, we repeated an identical permutation procedure for $\theta_{\mathrm{SC}_2}$ with $\Delta_{\mathrm{SC,intra}}$ as the target (in place of age), and found the observed association ($\rho = -0.96$) dwelling far beyond its own null band ($p < 1/500$). This finding provides further support for $\theta_{\mathrm{SC}_2}$'s lack of age-association reflecting the invariance of $\Delta_{\mathrm{SC,intra}}$ towards age in contrast to an identifiability failure.

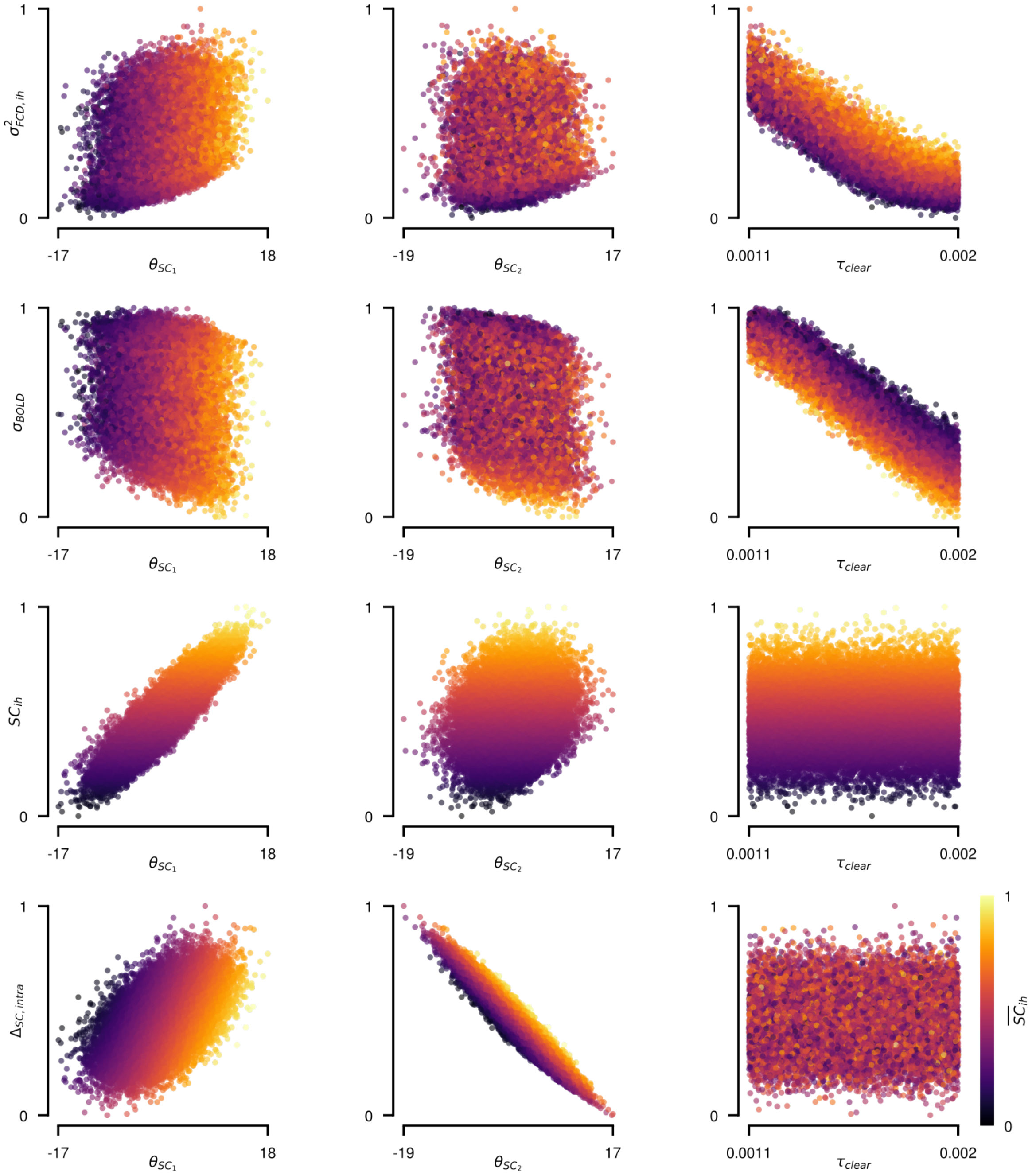

**Fig. S3**: **Full sensitivity map between the four training features and the three inferred parameters.** Each row shows one input feature ($\sigma^2_{\mathrm{FCD_{ih}}}$, $\sigma_{\mathrm{BOLD}}$, $\overline{SC}_{\mathrm{ih}}$, and $\Delta_{\mathrm{SC,intra}}$, top to bottom) plotted against all three inferred parameters across the $N$ = 20,000 synthetic training simulations, colour-coded by $\overline{SC}_{\mathrm{ih}}$. Columns correspond to $\theta_{\mathrm{SC}_1}$, $\theta_{\mathrm{SC}_2}$, and $\tau_{\mathrm{clear}}$, respectively.

#### S2. CrossCoder architecture selection across latent dimensionalities

The two-dimensional latent space $\boldsymbol{\theta}_{\mathrm{SC}} = (\theta_{\mathrm{SC}_1}, \theta_{\mathrm{SC}_2})$ employed throughout this study was chosen from a narrow sweep over candidate CrossCoder dimensionalities $n_{\mathrm{lat}} \in \{2, 3, 4\}$ (depicted in Methods, Section 1.2). This subsection accounts for the evidence supporting said choice, namely training/test reconstruction-MSE trace, held-out subject-identifiability confusion matrix, and pooled empirical-vs-reconstructed SC edge-weight fidelity (Fig. S2). As evident, all three 'architectures' coalesce to a stable train/test loss with no observable indication of overfitting, and complete held-out subject identifiability (100%). Our training demonstrated that reconstruction fidelity only marginally varies across latent space dimensionalities (Spearman $\rho = 0.90$ at $n_{\mathrm{lat}} = 2$ versus $\rho = 0.91$ at $n_{\mathrm{lat}} = 3$ and $n_{\mathrm{lat}} = 4$), hence inspiring our election of $n_{\mathrm{lat}} = 2$.

#### S3. Synthetic parameter-feature mappings

As briefly shown in main-text Fig. 4$a$, ADAM's structural latents $\boldsymbol{\theta}_{\mathrm{SC}}$ and its dopaminergic clearance timescale $\tau_{\mathrm{clear}}$ occupy largely complementary axes within the summary-statistics landscape. Supplementary Fig. S3 furnishes an exhaustive characterization of feature-parameter mappings across the full $N_{\mathrm{sim}}$ = 20,000 training simulations where synthetic features are coloured by $\overline{SC}_{\mathrm{ih}}$ in order to expose structural degradation (and thereby pseudo-personalized feature curves) as a third axis. We observed that $\theta_{\mathrm{SC}_1}$ and $\theta_{\mathrm{SC}_2}$ associate robustly and near-monotonically with their representative features, and only modestly with their contralateral structural feature ($r = 0.89$ vs. $r = 0.52$ for $\theta_{\mathrm{SC}_1}$, and $r = 0.22$ vs. $r = -0.95$ for $\theta_{\mathrm{SC}_2}$ with respect to $\overline{SC}_{\mathrm{ih}}$ & $\Delta_{\mathrm{SC,intra}}$), whereas both structural parameters bore a markedly weaker association with the functional summary statistics $\sigma^2_{\mathrm{FCD_{ih}}}$ and $\sigma_{\mathrm{BOLD}}$ ($|r| \leq 0.28$), a magnitude in keeping with that typically observed for structure–function correspondence (Supplementary Fig. S3).
In contrast, $\tau_{\mathrm{clear}}$ manifests a strong association with the BOLD data features $\sigma^2_{\mathrm{FCD_{ih}}}$ and $\sigma_{\mathrm{BOLD}}$ ($r = -0.91$ and $r = -0.95$, respectively) while remaining comparatively weak and diffuse with respect to the structural ones ($r = 0.00$ for $\overline{SC}_{\mathrm{ih}}$ and $r = -0.01$ for $\Delta_{\mathrm{SC,intra}}$; Supplementary Fig. S3). Taken together, our four-dimensional feature array establishes approximately orthogonal mappings across the structural and neuromodulatory axes, which constitutes the mechanistic prerequisite for our amortized estimator's capacity to disentangle structural and dopaminergic influences on the purported empirical observations.

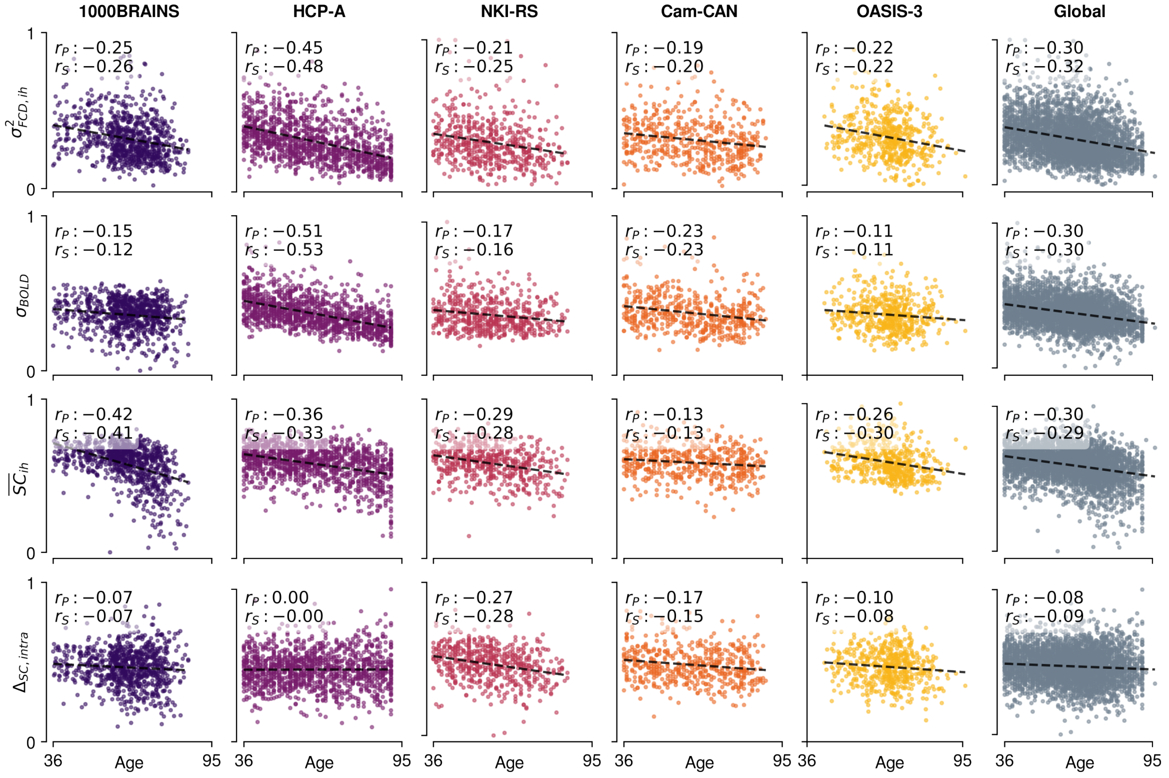


**Fig. S4**: **Chosen input features versus age, per cohort.** Rows show the four features entering the observation model ($\sigma^2_{\mathrm{FCD_{ih}}}$, $\sigma_{\mathrm{BOLD}}$, $\overline{SC}_{\mathrm{ih}}$, and $\Delta_{\mathrm{SC,intra}}$, top to bottom); columns show each of the five contributing cohorts (NKI-Rockland, Cam-CAN, HCP-Aging, OASIS-3, 1000BRAINS) followed by a pooled "Global" aggregate. Each panel reports Pearson ($r_P$) and Spearman ($r_S$) correlations with chronological age, with a linear regression fit (black, dashed).

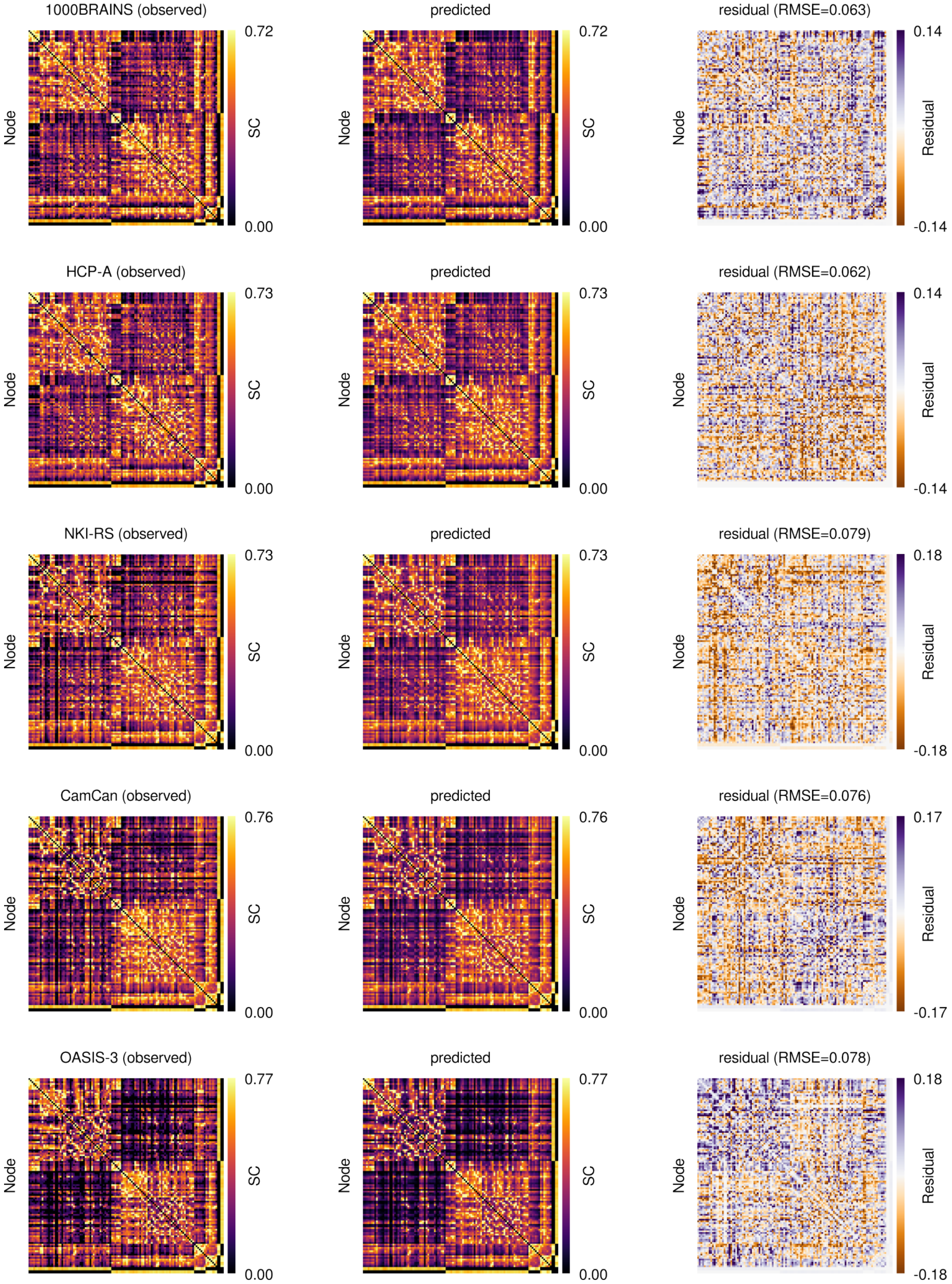


**Fig. S5**: **Subject-level structural connectome reconstruction fidelity, per cohort.** For one representative subject per cohort (**a,** 1000BRAINS; **b,** HCP-Aging/HCP-A; **c,** NKI-Rockland/NKI-RS; **d,** Cam-CAN; **e,** OASIS-3), observed structural connectivity matrix (left), CrossCoder-decoded reconstruction (middle), and their residual with the corresponding root-mean-square error (right).

### S4. Node-resolved simulations under a diffuse prior of clearance timescales

Simulating over a diffuse range of clearance timescales suggests an initial increase in fluidity, followed by a maximum, and its subsequent attenuation with further elevation in $\tau_{\text{clear}}$ (Main text Fig. 2*e*). However, this does not tell us of the exact node-level perturbations which result in those macroscale observations. We investigated this via simulations returning both the time-averaged neural state and BOLD dynamics (with biophysical constants, noise and integrator remaining unchanged), providing us with the D1 occupancy, firing rate, and rate variability of every node, here averaged over the 114 dopaminergically innervated nodes (in addition to fluidity; Supplementary Fig. S6). Besides corroborating the fluidity profile of Fig. 2*e* (Supplementary Fig. S6*a*), slowing clearance timescales over the abovementioned diffuse range initially enhances the structure–function coupling. Particularly, interhemispheric structure–function coupling rises from 0.03–0.09 to a maximum of 0.15–0.26, followed by an attenuation to 0.05–0.09 (Supplementary Fig. S6*c*). We restricted this correlation to interhemispheric edges, since attenuating only those weights would otherwise inflate a correlation pooled over all edges, and used a rank correlation, which is insensitive to the uniform rescaling of those weights. Note the concomitant changes in this coupling and fluidity (Supplementary Fig. S6*a*, *c*). Fluidity, coupling and rate variability all peak at $\tau_{\mathrm{clear}} \approx 10^{-3}$ at every structural level (Supplementary Fig. S6*a*, *c*, *e*), whereas the mean firing rate rises across the whole sweep (Supplementary Fig. S6*d*). Beyond this peak, fluidity falls as firing rate rises ($\rho = -0.99$ to $-1.00$ across structural levels; Supplementary Fig. S6*b*). Prolonged clearance may thus drive populations into a sustained high-activity regime in which their rates vary less over time, and interhemispheric fluidity declines with them.

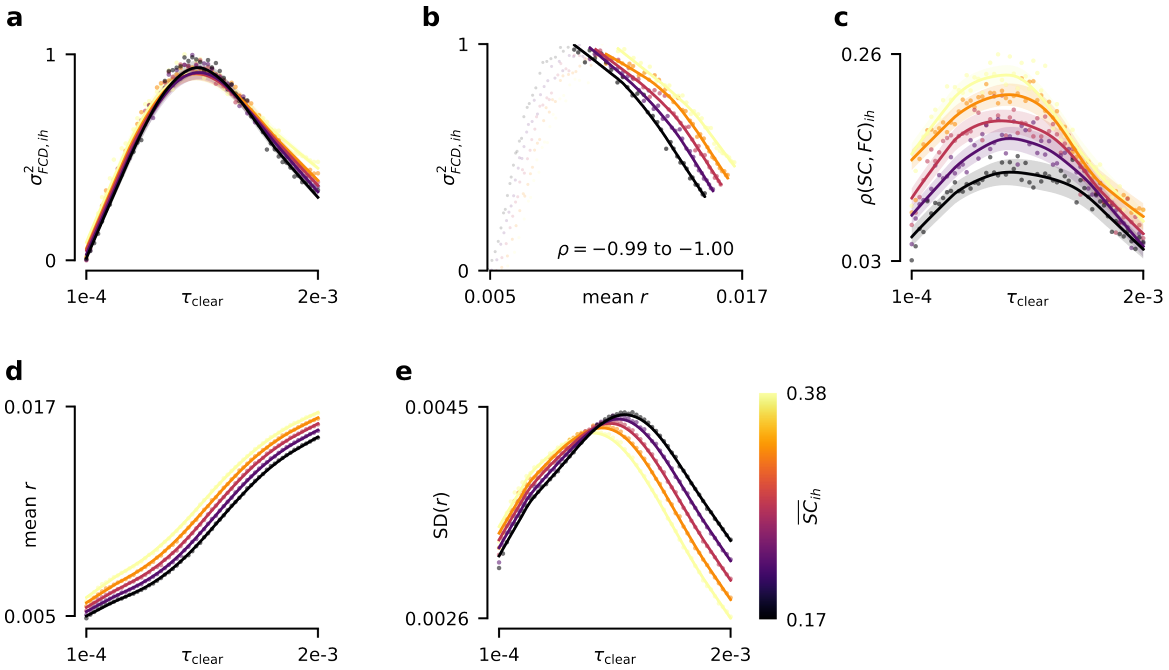


**Fig. S6**: **Lengthening dopaminergic clearance first strengthens interhemispheric structure–function coupling, then drives populations towards a saturated regime where coupling and fluidity fall together.** $\tau_{\mathrm{clear}}$ swept over $[1, 20] \times 10^{-4}$ (60 values) at five levels of interhemispheric attrition, obtained by attenuating the interhemispheric structural edges of one reference connectome (114-node block, as in Fig. 1*e*) and colour-coded by the resulting $\overline{SC}_{\mathrm{ih}}$; intrahemispheric edges are left intact, as is the separate dopaminergic-projection connectivity through which the DA sources drive every node. Every $(\tau_{\mathrm{clear}}, \overline{SC}_{\mathrm{ih}})$ pair is averaged over 10 independent random seeds. Lines are a locally weighted (LOWESS) trend through the points; $\sigma^2_{\mathrm{FCD_{ih}}}$ is min–max normalized across the sweep, firing rate and coupling are left in their own units. **a,** Interhemispheric fluidity $\sigma^2_{\mathrm{FCD_{ih}}}$. **b,** $\sigma^2_{\mathrm{FCD_{ih}}}$ against mean rate, restricted to the falling branch (Spearman $\rho$ per structural level); faint points trace the full sweep. **c,** Interhemispheric structure–function coupling: Spearman rank correlation between SC and simulated BOLD functional connectivity over interhemispheric edges. **d,** Mean population firing rate across the 114 dopaminergically innervated nodes, from the time-averaged neural state rather than BOLD. **e,** Mean node-level temporal SD of that rate.

#### S5. Empirical summary statistics versus age

Fig. S4 delineates both a per-cohort and a 'Global' portrait of the empirical age-dependent trajectories of our chosen summary statistics, which was briefly alluded to in main-text Fig. 1*h*–*j*. Table S1 reports the corresponding age associations across cohorts and the pooled aggregate. $\Delta_{\mathrm{SC,intra}}$ shows no significant age-related decline in 1000BRAINS and HCP-A, which are sourced and preprocessed via their own externally released pipelines, whereas a decline is present in two of the three in-house-processed cohorts, NKI-RS and Cam-CAN. This contrast may stem from a specific parameter choice within our unified in-house pipeline rather than a genuine biological effect (see Methods, Sections 1.1.1, 1.1.2, 1.1.3). Nevertheless, since $\theta_{\mathrm{SC_2}}$ encodes for inter-individual differences in the signed left–right difference in intrahemispheric SC ($\Delta_{\text{SC,intra}}$) rather than age-dependent degradation, the abovementioned slight age-related decline in NKI-RS/Cam-CAN bears no effect on our findings in terms of $\tau_{\text{clear}}$ or $\theta_{\text{SC}_1}$.

#### S6. Subject-level structural connectome reconstruction exemplars

We supplement the robust reconstruction fidelity of the CrossCoder with granular, subject-scale exemplars of the same metric. Supplementary Fig. S5 juxtaposes the observed, CrossCoder-decoded, and residual (observed minus decoded) SC matrices for one subject per cohort, with relatively low reconstruction RMSE across cohorts (1000BRAINS: 0.063; HCP-A: 0.062; NKI-RS: 0.079; Cam-CAN: 0.076; OASIS-3: 0.078, expressed in harmonized, log-transformed units), which provides further evidence for reliable encapsulation of individual-level structural nuances by the two-dimensional latent space $\boldsymbol{\theta}_{\text{SC}}$ parameters.

#### S7. Per-task validation of the cognitive composite across cohorts

The global cognitive composite Cog used throughout this study is derived separately for each cohort from its own battery of cognitive tasks (Methods, Section 1.1.5). Below, we delineate said battery, which is visualized individually in Supplementary Fig. S7. 18 measures were selected from the 1000BRAINS cohort: a problem-solving test, selective attention, Stroop interference, figural fluency, a processing-speed measure, a concept-shifting (Trail-Making-type) measure, figural memory, forward/backward visuospatial working-memory span, visual memory, forward/backward digit span, phonemic and semantic verbal fluency (including switching variants), and two verbal memory scores. 6 measures were selected from HCP-Aging (HCP-A): five NIH Toolbox Cognition instruments (Pattern Comparison Processing Speed, Dimensional Change Card Sort, Flanker Inhibitory Control and Attention, List Sorting Working Memory, Picture Sequence Memory), and the Montreal Cognitive Assessment (MoCA) total score. 14 measures were selected from Cam-CAN: the Cattell fluid-intelligence test, the Hotel multitasking test (deviation from the optimal allocation of time and number of sub-tasks attempted), unfamiliar-face matching, famous-face naming and emotional-picture recognition memory (Benton Faces, Famous Faces, Emotional Memory), visual short-term memory (capacity and precision), tip-of-the-tongue ratio and the number of famous faces named correctly, picture-naming accuracy on the unprimed baseline block, two semantic/syntactic-ambiguity reaction-time measures, and the coefficient of variation of in-scanner reaction time. Finally, 12 measures were selected from OASIS-3: Trail Making A and B, Digit Symbol substitution, Letter-Number Sequencing, a task-switching measure, Logical Memory (immediate and delayed recall), Selective Reminding Test free recall, forward/backward digit span, and animal/vegetable category fluency. We did not manage to obtain the phenotypic data for the NKI-Rockland Sample in time for the conclusion of this analysis, and as such, cognitive performance measures for this cohort are not included in the computation of Cog.

**Table S1**: **Age associations of the four input features, per cohort.** Pearson ($r_P$) and Spearman ($r_S$) correlations between each summary statistic and age, reported for every contributing cohort and for the "Global" aggregate. Values correspond to the regression fits shown in Fig. S4.

| | 1000BRAINS | | HCP-A | | NKI-RS | | Cam-CAN | | OASIS-3 | | Global | |
|---|---|---|---|---|---|---|---|---|---|---|---|---|
| Feature | $r_P$ | $r_S$ | $r_P$ | $r_S$ | $r_P$ | $r_S$ | $r_P$ | $r_S$ | $r_P$ | $r_S$ | $r_P$ | $r_S$ |
| $\sigma^2_{\text{FCD}_{\text{ih}}}$ | −0.25 | −0.26 | −0.45 | −0.48 | −0.21 | −0.25 | −0.19 | −0.20 | −0.22 | −0.22 | −0.30 | −0.32 |
| $\sigma_{\text{BOLD}}$ | −0.15 | −0.12 | −0.51 | −0.53 | −0.17 | −0.16 | −0.23 | −0.23 | −0.11 | −0.11 | −0.30 | −0.30 |
| $\overline{SC}_{\text{ih}}$ | −0.42 | −0.41 | −0.36 | −0.33 | −0.29 | −0.28 | −0.13 | −0.13 | −0.26 | −0.30 | −0.30 | −0.29 |
| $\Delta_{\text{SC,intra}}$ | −0.07 | −0.07 | +0.00 | −0.00 | −0.27 | −0.28 | −0.17 | −0.15 | −0.10 | −0.08 | −0.08 | −0.09 |

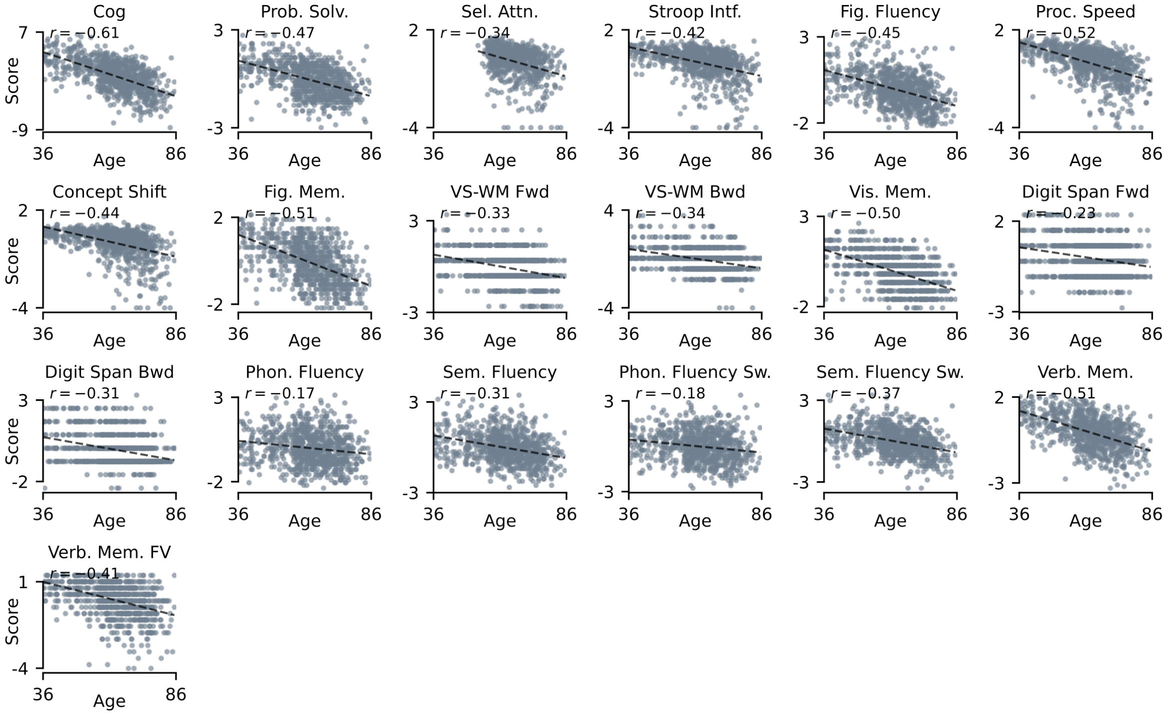


**Supplementary Fig. S7. Standardized, sign-aligned single-task cognitive performance versus age, per cohort.** Each panel shows one neuropsychological task's (z-scored) score against chronological age, with a linear regression fit and the corresponding Pearson $r$; the leading panel of each cohort shows the global composite Cog itself. **This page:** 1000BRAINS. HCP-A, Cam-CAN and OASIS-3 follow, one cohort per page.

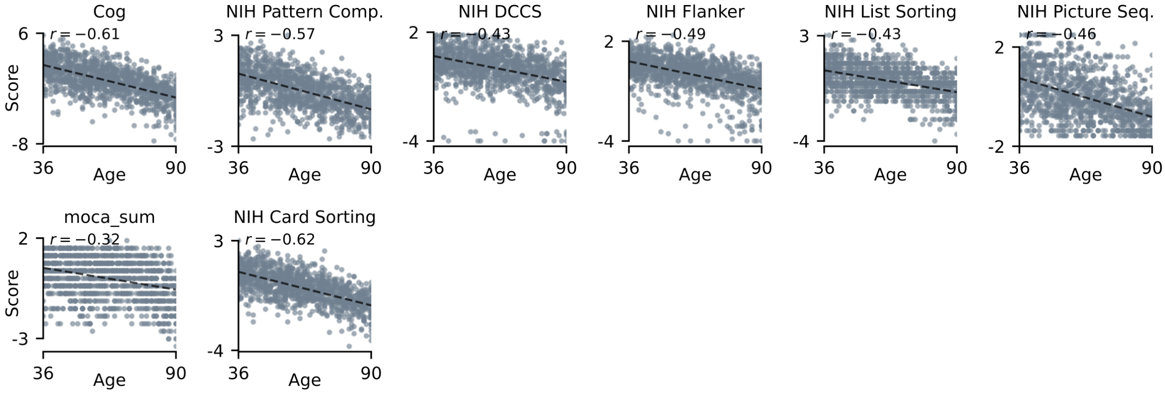


**Supplementary Fig. S7 (continued). HCP-Aging (HCP-A).**

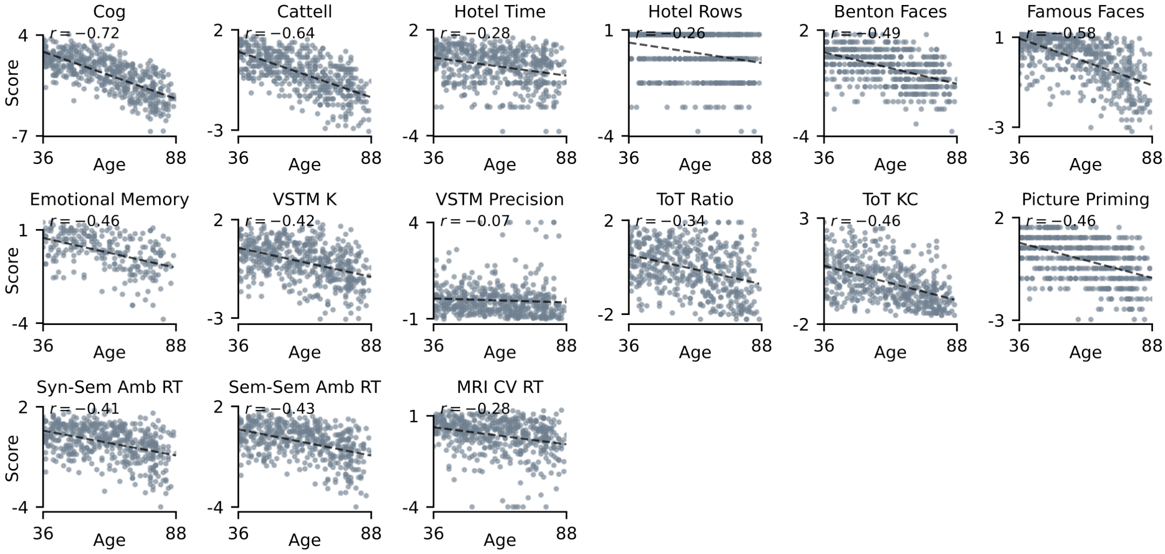


**Supplementary Fig. S7 (continued). Cam-CAN.**

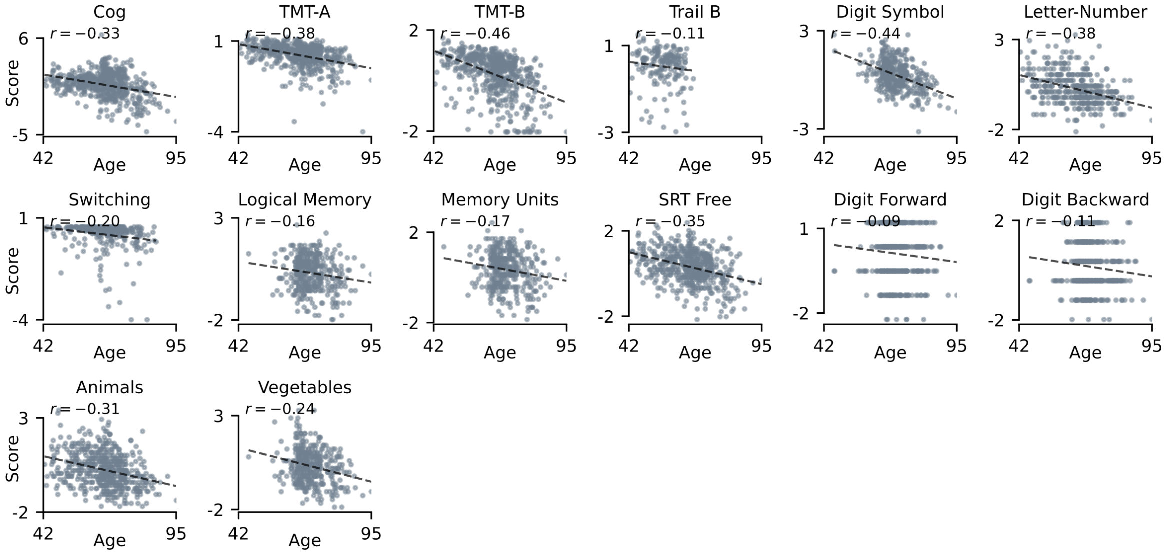


**Supplementary Fig. S7 (continued). OASIS-3.** NKI-Rockland cognitive phenotypes were unavailable for the present analysis.

As described in Methods, we sign-aligned all of the z-scored tasks via their empirical Spearman correlation with age (Methods, Section 1.1.5), subsequently holding the resulting directions fixed. Applying this procedure flagged Trail Making completion times, Stroop interference scores, the 1000BRAINS selective-attention time, the Cam-CAN Hotel-task time-allocation deviation, the tip-of-the-tongue ratio, the two sentence-ambiguity reaction times, and the coefficient of variation of in-scanner reaction time for inversion. The task-level coefficients reported below, and those in Supplementary Fig. S7, are Pearson correlations computed after this alignment, whereas the alignment itself proceeded on Spearman correlations. We report the standardized, sign-aligned task-level correlations for every individual cognitive measure with age, partitioned by the four cohorts with a complete task-level battery (1000BRAINS, HCP-A, Cam-CAN, and OASIS-3). In every cohort, the composite exhibited a significant decline with age ($r = -0.61$ in 1000BRAINS, $r = -0.60$ in HCP-A, $r = -0.72$ in Cam-CAN, and $r = -0.32$ in OASIS-3), with the substantial majority of constituent tasks recapitulating this decline individually across domains of processing speed, executive function, and memory. A small fraction of individual tasks exhibited weak or negligible age-association (e.g. forward digit span in OASIS-3, $r = -0.08$; and the Cam-CAN colour short-term-memory precision, $r = -0.07$), which may reflect task-specific sensitivity, range restriction, or non-linear age trajectories. The value of the composite delineated here lies in showing its concurrence with the vast majority of individual tasks, as opposed to being driven by a handful of dominant ones. The pooled composite hence serves as a reliable global index of age-related cognitive decline (Supplementary Fig. S7). The first component dominates its spectrum in every cohort, accounting for 39.7%, 54.3%, 31.8% and

43.1% of task-level variance in 1000BRAINS, HCP-A, Cam-CAN and OASIS-3 respectively, against a second component of 9.2%, 13.8%, 13.2% and 9.9%.

#### S8. Disentangling the influence of structural decay versus dopamine clearance timescales on cognitive performance

The moderation analysis portrayed in main-text Fig. 4*g*–*i* is augmented by a nested-regression, aiming to quantify how much of global cognitive composite Cog variance is explained through structural degradation ($\overline{SC}_{\mathrm{ih}}$), and how much of the unexplained variance left by $\overline{SC}_{\mathrm{ih}}$ may be further explained by $\tau_{\mathrm{clear}}$. We therefore fit three ordinary-least-squares models on raw (non-residualized) Cog, with every model bearing a quadratic age term as a covariate. In contrast with the interaction model shown in main-text Fig. 4*g*, *h*, we deliberately did not regress age out beforehand, as this query requires the age-explained variance of Cog to remain accessible for $\overline{SC}_{\mathrm{ih}}/\tau_{\mathrm{clear}}$ to compete for: $M_0: \mathrm{Cog} \sim \mathrm{age}_c + \mathrm{age}_c^2$; $M_1: M_0 + \overline{SC}_{\mathrm{ih}}$; $M_2: M_1 + \tau_{\mathrm{clear}}$. Here, we measured incremental $R^2$ relative to $M_0$ for both $M_1$ and $M_2$, computed the partial $R^2$ of $\tau_{\mathrm{clear}}$ over $M_1$ as $(R^2(M_2) - R^2(M_1))/(1 - R^2(M_1))$ (the fraction of variance left unexplained by $M_1$ that $\tau_{\mathrm{clear}}$ accounts for), and $F/p$ from the nested $F$-test against the immediately preceding model (Table S2, with the full coefficients in Table S3). We observed that for subjects with 'exhaustive data' ($n = 3{,}106$, i.e. subjects with complete Cog, $\overline{SC}_{\mathrm{ih}}$, $\tau_{\mathrm{clear}}$, and age data), $\overline{SC}_{\mathrm{ih}}$ explained a modest but highly significant increment of variance over age alone (incremental $R^2 = 0.010$, $F(1, 3102) = 45.0$, $p = 2.4 \times 10^{-11}$). Moreover, $\tau_{\mathrm{clear}}$ (entered as an unconditional covariate on top of $M_1$) added negligible additional variance (partial $R^2 = 0.00002$, $F(1, 3101) = 0.07$, $p = 0.79$). This negligible unconditional contribution of $\tau_{\mathrm{clear}}$ corroborates the findings of the moderation analysis depicted in main-text Fig. 4*g*–*i*, as the cognitive pertinence of DA clearance timescales is an interaction jointly gated by structural deterioration and age, rather than a global main effect. As such, an additive model which aggregates all SC and age strata together is expected to average out this gated contribution towards zero. Finally, restricting these three models to the 56–76-year age band previously identified by the moderation analysis (in the absence of degradation-stratification) corroborates the above ($n = 1{,}590$; SC incremental $R^2 = 0.023$, $p = 3.9 \times 10^{-10}$; $\tau_{\mathrm{clear}}$ partial $R^2 = 0.002$, $p = 0.061$). Here, structural integrity carries the explanatory variance, whereas the unconditional contribution of $\tau_{\mathrm{clear}}$ remains below significance even within the band in which the interaction is strongest (due to the absence of an explicit $\tau_{\mathrm{clear}} \times \overline{SC}_{\mathrm{ih}}$ term). Taken together, these additive analyses locate the cognitive variance in structural integrity rather than in $\tau_{\mathrm{clear}}$ as a main effect. The present additive models however, contain no $\tau_{\mathrm{clear}} \times \overline{SC}_{\mathrm{ih}}$ interaction term, and thus cannot identify whether this association hinges on structural decay (rather than age alone). We therefore augment the findings established here with the results of the explicit interaction test illustrated in main-text Fig. 4*g*–*i* (Methods, Section 1.5).

**Table S2**: Nested-OLS variance decomposition of the cognitive composite Cog

| Model | $R^2$ | Incremental $R^2$ | Partial $R^2$ | $F$ | $p$ |
|---|---|---|---|---|---|
| $M_0$: $\mathrm{Cog} \sim \mathrm{age}_c + \mathrm{age}_c^2$ | 0.313 | – | – | – | – |
| $M_1$: $M_0 + \overline{SC}_{\mathrm{ih}}$ | 0.323 | 0.010 | – | 44.98 | $2.4 \times 10^{-11}$ |
| $M_2$: $M_1 + \tau_{\mathrm{clear}}$ | 0.323 | 0.010 | 0.00002 | 0.07 | 0.79 |

**Table S3**: Full nested-OLS coefficients underlying Table S2

| Model | Term | Estimate | SE | $t$ | $p$ |
|---|---|---|---|---|---|
| $M_0$ | Intercept | 0.195 | 0.042 | 4.63 | $3.7 \times 10^{-6}$ |
| $M_0$ | $\text{age}_c$ | −0.087 | 0.002 | −37.56 | $< 10^{-200}$ |
| $M_0$ | $\text{age}_c^2$ | −0.0009 | 0.0002 | −5.94 | $3.1 \times 10^{-9}$ |
| $M_1$ | Intercept | −1.011 | 0.185 | −5.47 | $4.7 \times 10^{-8}$ |
| $M_1$ | $\text{age}_c$ | −0.082 | 0.002 | −33.87 | $< 10^{-200}$ |
| $M_1$ | $\text{age}_c^2$ | −0.0008 | 0.0002 | −5.11 | $3.4 \times 10^{-7}$ |
| $M_1$ | $\overline{SC}_{\text{ih}}$ | 1.990 | 0.297 | 6.71 | $2.4 \times 10^{-11}$ |
| $M_2$ | Intercept | −0.953 | 0.286 | −3.33 | 0.0009 |
| $M_2$ | $\text{age}_c$ | −0.082 | 0.003 | −32.16 | $< 10^{-195}$ |
| $M_2$ | $\text{age}_c^2$ | −0.0008 | 0.0002 | −5.11 | $3.4 \times 10^{-7}$ |
| $M_2$ | $\overline{SC}_{\text{ih}}$ | 1.963 | 0.314 | 6.24 | $4.8 \times 10^{-10}$ |
| $M_2$ | $\tau_{\text{clear}}$ | −0.074 | 0.283 | −0.26 | 0.792 |

### S9. Cognitive tasks grouped by domain

We regroup the same battery of cognitive measures shown in Methods, Section 1.1.5 by a canonical cognitive-domain taxonomy to allow for a clear juxtaposition of cognitive domain across cohorts (NKI-Rockland is omitted as previously stated, due to its cognitive data not being available for the present analysis). Our domain designation follows the NIH Toolbox Cognition Battery's official guideline for HCP-A tasks (Weintraub et al. 2013), standard neuropsychological classification for the classic tests shared across 1000BRAINS/OASIS-3 (Trail Making, Stroop, digit span, category fluency, figural/design fluency; (Lezak et al. 2012)), and the Cam-CAN's own task descriptions for its cohort-specific instruments (Shafto et al. 2014). Domain and function assignments, test identities and description of each measure for 1000BRAINS were provided by the cohort's own investigators and correspond to the battery documented and tabulated in Caspers et al. (2014); Jockwitz et al. (2021). Forward span tasks are hence designated as short-term memory, whilst only the backward (manipulation) conditions are taken as working memory; said distinction is applied uniformly across the four cohorts. It should be noted that the domain grouping presented below serves a presentational purpose alone, and remains provisional, since several measures overlap across domains (such as the digit span or picture-priming tasks). Its sole utility lies in the comparison of domains across the cohorts' task batteries, whereas Cog itself is computed directly from the first PC of the unaggregated, unweighted, and sign-aligned task-scale values (Methods, Section 1.1.5). We further excluded the Montreal Cognitive Assessment (MoCA) from this aggregation, due to it being a composite rather than a single-construct measure, despite said test still entering the computation of Cog via HCP-A's task list. Table S4 juxtaposes domain coverage across the four cohorts at the level of test names.

**Table S4**: Domain-grouped cognitive tasks across cohorts

| Domain | 1000BRAINS | HCP-A | Cam-CAN | OASIS-3 |
|---|---|---|---|---|
| Attention | Alters-Konzentrations-Test | – | – | – |
| Processing speed | Trail Making Test A | NIHTB Pattern Comparison | In-scanner simple reaction time (CV) | Trail Making Test A; WAIS-R Digit Symbol |
| Executive function | Farb-Wort-Interferenztest; Trail Making B−A; Fünf-Punkte-Test | NIHTB Dimensional Change Card Sort; NIHTB Flanker | Hotel multitasking test (time-allocation deviation; tasks attempted) | Trail Making Test B; task-switching accuracy |
| Fluid reasoning | Leistungsprüfsystem 50+, Subtest 3 | – | Cattell Culture Fair, Scale 2 | – |
| Short-term memory | Zahlennachsprechen, forward; Block-Tapping-Test, forward; Visual pattern | – | Colour visual short-term memory (capacity, precision) | WMS Digit Span forward |
| Working memory | Zahlennachsprechen, backward; Block-Tapping-Test, backward | NIHTB List Sorting | – | WMS-III Letter-Number Sequencing; WMS Digit Span backward |
| Episodic memory | Verbaler Gedächtnistest (learning, delayed recall); Benton-Test | NIHTB Picture Sequence Memory | Emotional Memory (object recognition) | WMS Logical Memory IA and IIA; Selective Reminding Test, free recall |
| Face and person knowledge | – | – | Benton Facial Recognition; Famous Faces | – |
| Language and verbal fluency | Regensburger Wortflüssigkeitstest (phonemic, semantic, phonemic switching, semantic switching) | – | Tip-of-the-tongue task; picture naming; sentence ambiguity resolution | Category fluency (animals, vegetables) |
| Global screening | – | Montreal Cognitive Assessment | – | – |

### References


Caspers S, Moebus S, Lux S, et al (2014) Studying variability in human brain aging in a population-based German cohort—rationale and design of 1000BRAINS. Frontiers in Aging Neuroscience 6:149. doi: 10.3389/fnagi.2014.00149

Jockwitz C, Wiersch L, Stumme J, et al (2021) Cognitive profiles in older males and females. Scientific Reports 11(1):6524. doi: 10.1038/s41598-021-84134-8

Lezak MD, Howieson DB, Bigler ED, et al (2012) Neuropsychological Assessment, 5th edn. Oxford University Press, New York

Shafto MA, Tyler LK, Dixon M, et al (2014) The Cambridge Centre for Ageing and Neuroscience (Cam-CAN) study protocol: a cross-sectional, lifespan, multidisciplinary examination of healthy cognitive ageing. BMC Neurology 14:204. doi: 10.1186/s12883-014-0204-1

Weintraub S, Dikmen SS, Heaton RK, et al (2013) Cognition assessment using the NIH Toolbox. Neurology 80(11 Suppl 3):S54–S64. doi: 10.1212/WNL.0b013e3182872ded
